# Spring-loaded DNA origami arrays as energy-supplied hardware for modular nanorobots

**DOI:** 10.1101/2024.09.30.615428

**Authors:** Martina Pfeiffer, Fiona Cole, Dongfang Wang, Yonggang Ke, Philip Tinnefeld

## Abstract

DNA origami nanodevices allow us to mimic cellular functions in a rationally controlled manner. They describe machineries which respond to environmental stimuli by conducting different tasks. To date, this mostly is achieved by constructing conformational two-state switches which upon activation by stimuli change their conformation resulting in the performance of a priorly programmed task. Their applicability however is often limited to a single, specific stimuli – output combination due to their intrinsic properties as two-state systems only. This makes expanding them further to include multiple stimuli/ outputs challenging. Here, we address this problem by introducing reconfigurable DNA origami arrays as a coupled network of two-state systems. We use this network to create a universal design strategy in which different operational units can be incorporated into any two-state system of our nanorobot. The resulting nanorobot is capable of receiving different stimuli, computing the response to the received stimuli using multi-level Boolean logic gating and yielding multiple programmed output operations with controlled order, timing and spatial position. We expect that this strategy will be a crucial step towards further developing DNA origami nanorobots for applications in various technological fields.

## Introduction

Over the last decades, the DNA origami technique^1,2^ has emerged as an indispensable tool for designing devices capable of emulating functions and properties of naturally occurring machines on the nanoscale and beyond that, increasingly perform robotic tasks such as sensing, computing and actuating.^3–8^

DNA origami involves the folding of a long single-stranded scaffold strand into a custom shape by up to hundreds of oligonucleotide “staple” strands. Most current DNA origami nanodevices are designed and optimized to perform a specific operation such as cargo release^4,7–9^, a rotational motion^3,10,11^ or a chemical reaction^5,12^ after sensing chemical or physical stimuli^13^. This is often achieved by inducing a single, relatively simple conformational change in the nanodevice, causing it to act as a two-state switch whose operation may optionally include simple AND or OR^6,7^ gate logics. The conformational change alters the proximity of interacting players of the operational parts of the nanodevice, resulting in the performance of a defined operation. However, the fact that this concept is based on a single conformational change makes expanding it further challenging. For the realization of more sophisticated nanodevices capable of autonomously performing a series of operations in response to different combinations of environmental stimuli, so far no general concept exists.

Here, we present the DNA origami nanorobot platform SEPP (for **S**erial **E**xecution of **P**rogrammable **P**rocesses) that uses a reconfigurable DNA origami array system composed of multiple, structurally similar blocks – so-called antijunctions as basis for multistep operations (Fig. 1a,b).^14–16^ Antijunctions are small symmetric constructs containing four DNA duplex domains of equal length which are pairwise stacked as well as four dynamic nicking points. They exist in two stable conformations with reversed stacking order between which they can switch via an instable open conformation.^14^ In reconfigurable DNA origami arrays, multiple antijunctions are coupled to each other by the scaffold strand which threads through the whole system. It interconnects the individual antijunctions and forces them to all adapt the same conformation. Induced by the hybridization of fuel DNA strands to certain antijunctions at the edge of the structure, the conformation of the whole system can be reconfigured in a diagonal stepwise manner (Fig. 1b). In each of the steps, a row of antijunctions in the system undergoes a conformational change, ultimately resulting in the reconfiguration of the whole structure. Full reconfiguration of DNA origami arrays generally was used to activate different proximity-induced operations^17,18^ such as the onset of catalytic activity^19^, the performance of different pattern operations involving writing, erasing and shifting^20^, the generation of an optical output signal^17^, or the release of cargo DNA strands^21^. However, these stimuli did not target the conformation of a specific antijunction in the system. They rather targeted the overall conformation of the system itself disregarding the potential of its intermediates and reducing it to a simple two-state switch.

**Figure 1:**
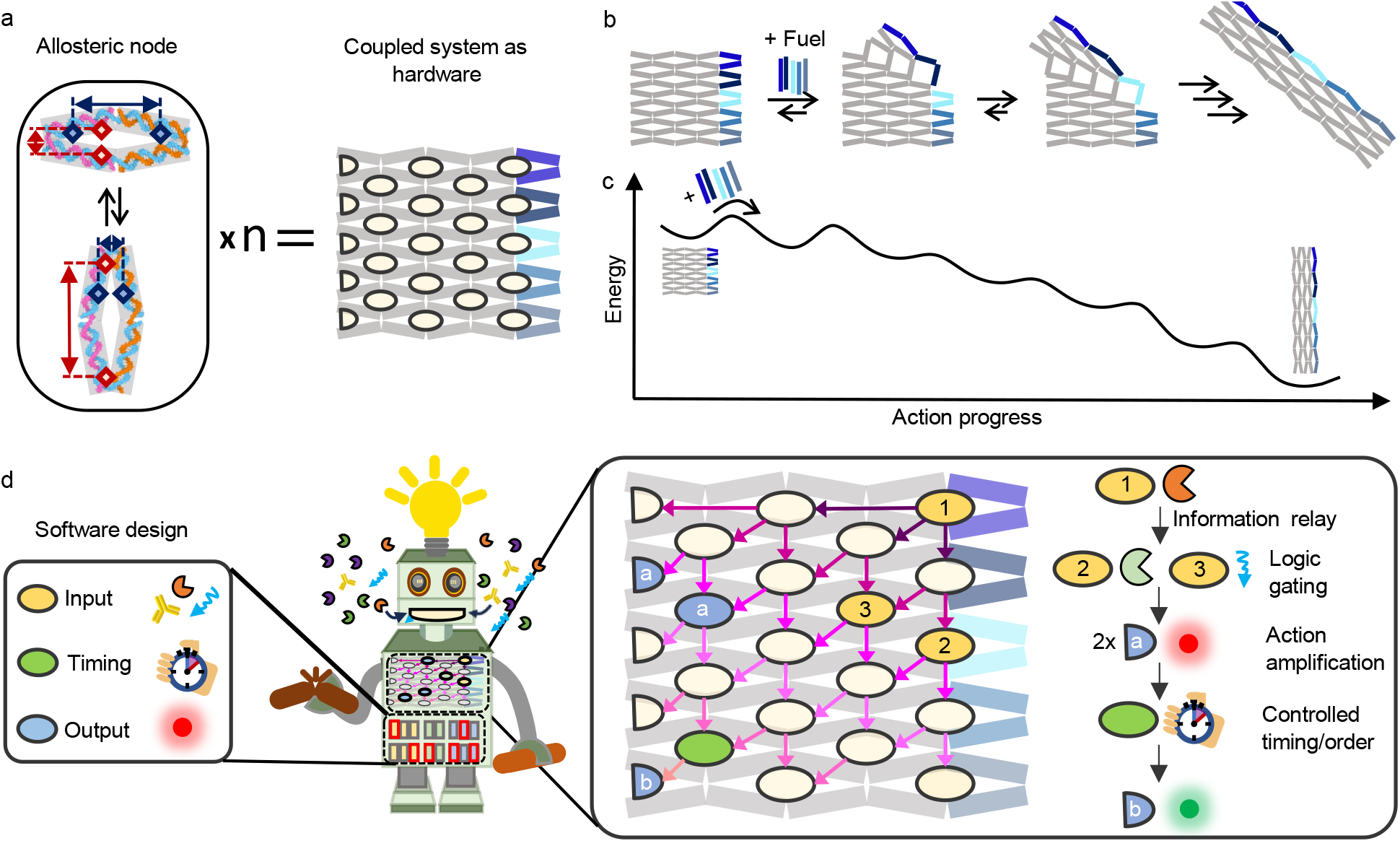
Concept of a modular reconfigurable DNA origami nanorobot. (a) Sketch of multiple allosteric nodes coupled together in a reconfigurable DNA origami array forming the hardware of the design of the nanorobot. The nodes are indicated by ellipses at the corresponding positions in the DNA origami array. The basis unit of an allosteric node is a so-called anti-junction which switches between two conformational states (left). Due to the symmetric nature of the antijunction, the switching can be used to both increase (red) and decrease (dark blue) the distance between markers placed on the antijunction depending on which domains they are placed on. (b) Sketch of the reconfiguration process of the unmodified DNA origami array structure upon hybridization of fuel DNA strands (blue lines) to the right side of the structure. (c) Simplified sketch of the energy landscape of the multistep reconfiguration process.^21^ (d) The software of the designed nanorobot is formed by developing generalized strategies to encode environmental responsiveness to different target, timing, and output operations into single allosteric nodes (left panel). The nanorobot is then formed by programming different nodes of the DNA origami array hardware with the software (middle panel). The arrows hereby represent the connectivity between the different units given by the energy landscape of the hardware. This enables the robot to respond autonomously to targets in its environment in a pre-programmed, multistep manner (right panel).

In contrast, SEPP interprets reconfigurable DNA origami arrays as two-dimensional networks of coupled two-state switches represented by their individual antijunctions. SEPP builds on previous studies that uncovered the energy landscape of the reconfiguration process (Fig. 1c)^14,15,21^ and showed how – by incorporation of energy barriers – the coupling between different antijunctions is rationally altered to retard or even fully stop the reconfiguration at a specified intermediate step.^21^ In our approach, a reconfigurable DNA origami array constitutes the “hardware” of a DNA nanorobot which can be programmed by a software framework described in this work (Fig. 1d).

With its individual antijunctions, the hardware provides a set of nodes with known connectivity given by the network’s energy landscape as well as a defined starting point. With the fuel DNA strands, the hardware also provides an energy source to create a spring-loaded allosteric driving force for the nanorobot’s autonomous action. In our software framework, each antijunction within the system is considered as a potential input, output or timing node. By subscribing them with a functionality, the energy landscape of the transformation process is altered at the position where the corresponding node changes its conformation.

The universal and rational approach of the combined hardware and software package is demonstrated with different inputs, such as enzyme activities, proteins, DNA and light. The inputs are modularly combined with a range of operations, including cargo release, fluorescence on- and offset and signal amplification to achieve arbitrary input – output combinations on demand (Fig. 1d). Additionally, we highlight how our understanding of both the pathway and energy landscape of the reconfiguration process enables us to implement order dependencies, timing control and multi-level logic gating as well as simplicity of designing antagonistic operations by exploiting the antijunction’s symmetric nature (Fig. 1a). As such, this merge of hardware and software promises a new era of versatile nanoscale devices.

## Results

SEPP is based on a small reconfigurable DNA origami array structure composed of 5 x 2.5 antijunctions^14^ that can be transformed by hybridizing five fuel DNA strands to the right side of the structure (Fig. 1a, b).^21^ The transformation process is characterized by five intermediates (Fig. 1c). Our software framework targets the energy barriers between these intermediates and the start and the end point of the transformation. To characterize SEPP at the single-molecule level with fluorescence microscopy, we additionally incorporate dye-quencher pairs for reporting on the state of individual antijunctions and a biotinylated ssDNA for surface-immobilization on BSA-biotin/Neutravidin coated cover slips (Supplementary Fig. 1).

### Design of environmental-responsive input nodes

Environmental responsiveness is encoded into the antijunction nodes by the introduction of locking units which stabilize one conformation of the targeted node over the other. This results in an energetic bias either hindering or favoring the transformation of the whole array (Fig. 2a). By varying the number of added fuel DNA strands, we spring-load the array system with varying degrees of tension and adjust the energy levels of the untransformed and transformed state of the array to lie close together such that presence of (multiple) locking units becomes the decisive factor for the transformation to occur. In presence of the corresponding environmental input, the units are unlocked and the energetic bias is removed, resulting in the release of the stored tension and an activation or inhibition of the transformation process, respectively. Based on this principle, we design locking units responsive to single-stranded DNA (ssDNA), restriction enzyme activity, light and antibodies (Fig. 2, Supplementary Fig. 2-4).

**Figure 2.**
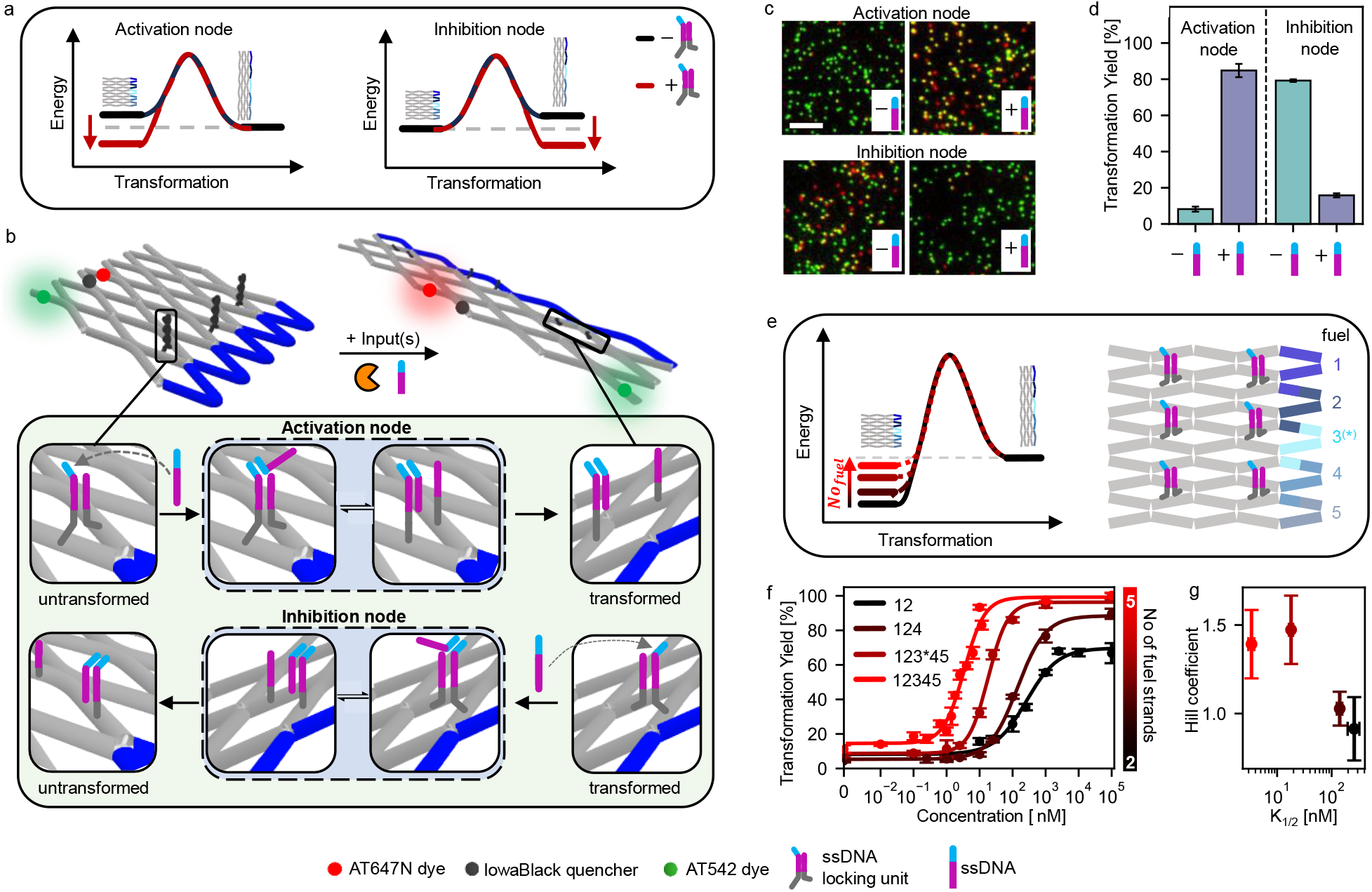
Environmentally responsive activation and inhibition nodes. (a) Energy schemes showing the influence of a locking unit in activation and inhibition nodes on the transformation process. In presence of their inputs, this effect is nullified. The energy schemes in absence of the locking units are programmed priorly by adding different combinations of fuel DNA strands. (b) Sketch of the DNA origami nanorobot bearing environmentally responsive nodes, a red dye-quencher FRET pair (ATTO647N-IowaBlackRQ) reporting on the state of the nanorobot and a green dye (ATTO542) for co-localization (upper panel). Nodes responsive to ssDNA activating and inhibiting the transformation are based on a reversible dsDNA lock which is unlocked via a toehold-mediated strand displacement reaction with a ssDNA input (lower panel). (c) Exemplary TIRF images of DNA origami nanorobots bearing ssDNA-responsive activation and inhibition nodes after incubation with fuel DNA strands 1,2 and 1,3* respectively and without and with the ssDNA input. (d) Corresponding transformation yields. (e) Strategy for tuning the responsive concentration window of the ssDNA input. (left) Increasing the number of added fuel DNA strands increases the tension in the spring-loaded system, resulting in a destabilization of the untransformed state of the array. (right) Employed DNA array system with six ssDNA locking units incorporated. The hybridization positions of fuel strands 1-5 are marked with color. Fuel strand 3* represents a shortened version of fuel strand 3. (f) Increasing the number of added fuel DNA strands from strands 1,2 over 1,2,4 and 1,2,3*,4,5 to 1,2,3,4,5, shifts the responsive window. (g) Impact of the number of added fuel DNA strands on K_1/2_ and the Hill coefficient. Error bars in (d,f) represent the standard deviation in the transformation yields calculated from at least three TIRF images. Error bars in (g) represent the fit error to the curves fitted in (f). Scalebar: 4 μm.

For ssDNA as an input, locking units are formed by two ssDNA strands protruding from two domains of the targeted antijunction nodes. The ssDNA strands contain a complementary section which hybridizes when both domains are in close proximity. Depending on which domains they are placed on, hybridization of the strands occurs either in the untransformed or transformed conformation of the nodes (Fig. 2b, upper and lower panel), resulting in their energetic stabilization. To enable unlocking, one of the protruding ssDNA strands is designed with a toehold overhang. This allows for unlocking by toehold-mediated strand displacement with a ssDNA input.

The successful implementation of the ssDNA locking units is verified using a fluorescence onset unit placed on an antijunction node transforming downstream in the transformation process. The fluorescence onset unit reports on the conformational state of the node it is placed on (Fig. 2b). It is based on a red dye-quencher probe (ATTO647N – IowaBlackRQ) positioned on two different domains of an antijunction node which are in close proximity in the untransformed state of the antijunction node. The increase in distance between these domains caused by the transformation of the corresponding node results in an increased fluorescence signal of the dye molecule. This allows to distinguish the untransformed from the transformed state of the node. We additionally incorporate an ATTO542 dye into the arrays and quantified the fraction of arrays in their transformed state (transformation yield) from dual-color single-molecule TIRF images of surface-immobilized array structures collected in absence and presence of the ssDNA input (Fig. 2c, Materials and Methods).

With a transformation yield of 10%, arrays with three ssDNA activation nodes incorporated and fuel DNA strands 1-2 are predominantly in their untransformed state in absence of ssDNA input. Upon addition of ssDNA input, the arrays transform, resulting in an increased transformation yield of 83%. In contrast, arrays with three ssDNA inhibition nodes incorporated and fuel DNA strand 1,3* show transformation yields of 80% and 16% in absence and presence of ssDNA input, demonstrating the antagonistic usability of the same DNA locking unit with near quantitative responses (Fig. 2d). As this process is reversible (Supplementary Fig. 5), our control over the energy landscape of the system provides additional access to tune the location (K_1/2_) and width of the responsive concentration window towards the ssDNA input without modifying the ssDNA locking unit itself. When incorporating six ssDNA activation units into the array, increasing the number of added fuel DNA strands from two to five leads to an increased mechanical strain exerted on the untransformed conformation of the array (Fig. 2e). This results in an over 75-fold decrease in K_1/2_ (from 260±60 nM to 3.4±0.4 nM) while simultaneously increasing the Hill coefficient from 0.9±0.2 to 1.4±0.2 and thus narrowing the responsive concentration window (Fig. 2f-g). Reducing the number of incorporated ssDNA activation units and replacing one of the ssDNA activation units with different units, introduces additional tuning strategies for the responsive window and allows shifting K_1/2_ down to 0.08±0.02 nM (Supplementary Fig. 6). In combination, all strategies allow shifting K_1/2_ over 3000-fold without modifying the ssDNA locking unit itself.^6^ This promises a good adaptability of these tuning strategies towards other inputs without the need for re-engineering the input-responsive locking units to create different affinities towards the inputs.

We then demonstrate the simple adaptability of our universal design approach towards other inputs. The designs of input nodes responsive to restriction enzyme activity and light are based on ssDNA locking units containing an enzyme-specific restriction site and a photocleavable linker, respectively. The locks are cleaved in presence of active restriction enzyme or light of a specified wavelength, effectively unlocking them (Supplementary Fig. 2, 3). When implemented as activation nodes in DNA array systems, they responded specifically to their respective inputs (XhoI, StuI and BamHI restriction enzyme and light of 365 nm) by transforming quantitatively only in their presence (Supplementary Fig. 2, 3) after addition of fuel DNA strands which can also be pre-loaded onto the structure (Supplementary Fig. 7). We further e tend PP’s spectrum of inputs to proteins which do not directly interact with DNA by designing an input unit responsive to IgG antibodies and demonstrating its application in inhibition nodes (Supplementary Fig. 4).

### Computation based on Boolean logics

In the design of our locking units, the presence of a single unit alone does not suffice to quantitatively initiate or inhibit the transformation process when the arrays are spring-loaded with all five fuel DNA strands (Supplementary Note 1, Supplementary Fig. 8). This design anticipates the rational creation of computation schemes, thus providing the ability to respond to diverse input combinations in a pre-programmed manner. We systematically characterize the impact of restriction enzyme locking units positioned at different nodes, both individually and in combinations (Fig. 3a, Supplementary Fig. 8).

**Figure 3.**
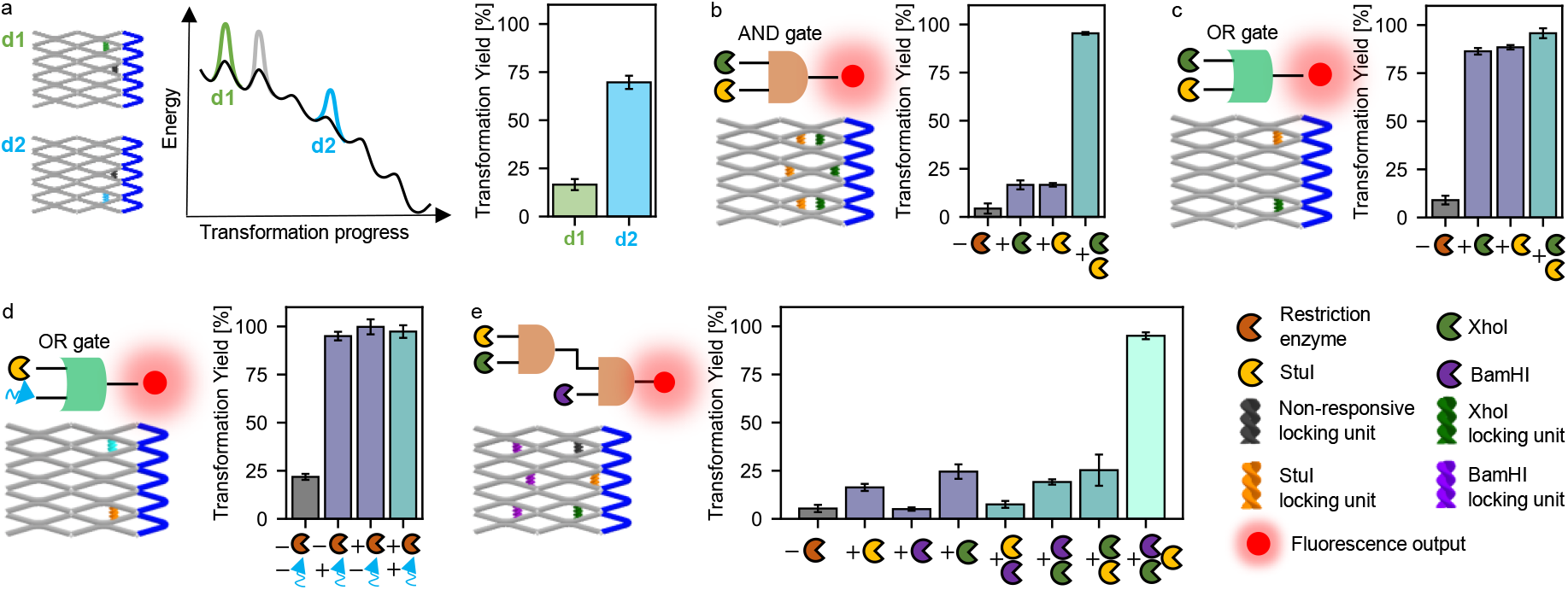
DNA origami array nanorobot processing one- and multi-level Boolean logic gates. One-level and multi-level Boolean logic gates responsive to combinations of restriction enzymes and light are implemented by combining different restriction enzyme/ light locking units and stabilization units. The molecular logic is programmed by the number and position of the incorporated units. Additionally, a red fluorescence onset unit is incorporated for reading out the state of the robot. (a) Comparison of the transformation yields of two array designs. The designs differ only in the position of one locking unit stabilizing the untransformed conformation marked in green (d1) and blue (d2) that target different positions in the energy landscape of the transformation process. The gray locking unit/ energy barrier is present in both designs. (b) Schematic design of a basic logic AND gate responsive to combinations of XhoI and StuI and obtained transformation yields. (c) Schematic design of a basic logic OR gate responsive to combinations of XhoI and StuI and obtained transformation yields. (d) Schematic design of a basic logic OR gate responsive to combinations of StuI and light and obtained transformation yields. (e) Schematic design of a 3xAND gate and obtained transformation yields. Error bars show the standard deviation from the mean. In addition to possible inputs, all DNA origami arrays are incubated with fuel DNA strands 1-5. Error bars represent the standard deviation in the transformation yields calculated from three TIRF images.

Notably, when comparing the effects of the same locking unit positioned at two different nodes in an otherwise identical design, we observe significant differences in the obtained transformation yields (Fig. 3a). When the locking unit is positioned at a node transforming during the first step of the transformation process, the transformation is inhibited. In contrast, placing the unit at a node transforming during the fourth step results in an increased transformation yield of 70%. This finding aligns with the profile of the energy landscape of the transformation process and is also consistent with all other studied locking unit combinations (Supplementary Note 1, Supplementary Fig. 8). The tilted profile of the energy landscape indicates a weakened effect of an incorporated locking unit the further downstream the corresponding antijunction node is in the transformation cascade. Using this dependence and placing multiple locking units specific to various inputs at predefined antijunction nodes on the same structure, we establish a computation framework to program responses to diverse input combinations based on Boolean logic gates.

We first implement basic Boolean AND and OR gates responsive to combinations of the restriction enzymes XhoI and StuI. We again use a fluorescence onset unit to confirm the designed responsiveness of the system. For the AND gate, only upon addition of both restriction enzymes a near quantitative transformation of all structures occurs (Fig. 3b, Supplementary Fig. 9) whereas for the OR gate already the addition of one restriction enzyme results in transformation yields of above 80% (Fig. 3c, Supplementary Fig. 9). The applicability of these gates is not limited to inputs of the same molecular class. Our modular design strategy of locking units for example allows the simple exchange of the XhoI locking unit with a light locking unit in the OR gate design while maintaining the functionality of the gate (Fig. 3d, Supplementary Fig. 9). This demonstrates how inputs of different molecular classes can be processed on the same structure in DNA origami arrays using SEPP.

Both the AND and the OR gate designs demonstrate low leakages in all logical FALSE conditions, with the TRUE state providing at least fourfold higher transformation yields in all cases. The low leakages of these basic gates allow expanding the concept further to also include multi-level logic gates. We use the multistep nature of the transformation cascade to – figuratively speaking – create cascades of multiple logic gates and connect them in series. Using the prior characterization of the effect of restriction enzyme locking units at different position as a basis, we design two multi-level logic gates of different complexity: A 3x AND gate which is comprised of two logic AND gates connected in series (Fig. 3e, Supplementary Fig. 10) as well as a gate which only gives a positive response if at least two of its three inputs are present (Supplementary Fig. 11). We find that both systems provide low leakages in all logical FALSE conditions that were statistically distinguishable from the transformation yields obtained for the TRUE states, demonstrating the successful implementation of the designed gates. Besides logic gating, the modularity offered by SEPP enables multiplexing as an additional pathway to process multiple inputs in parallel (Supplementary Fig. 12).

### Combining logical gated input operations with output operations

The aim of smart nanorobots not only includes programming responses to different input combinations but also the conduction of these responses if the required conditions are met. Beyond simple fluorescence onset units to generate output signals, we next demonstrate different output operations and combine input-responsive locking units with a more complex output operation. As an example, we choose a cargo release unit, which upon activation releases a fluorescently labelled (ATTO542) cargo DNA strand from the system (Fig. 4a, Supplementary Note 2).^21^

**Figure 4.**
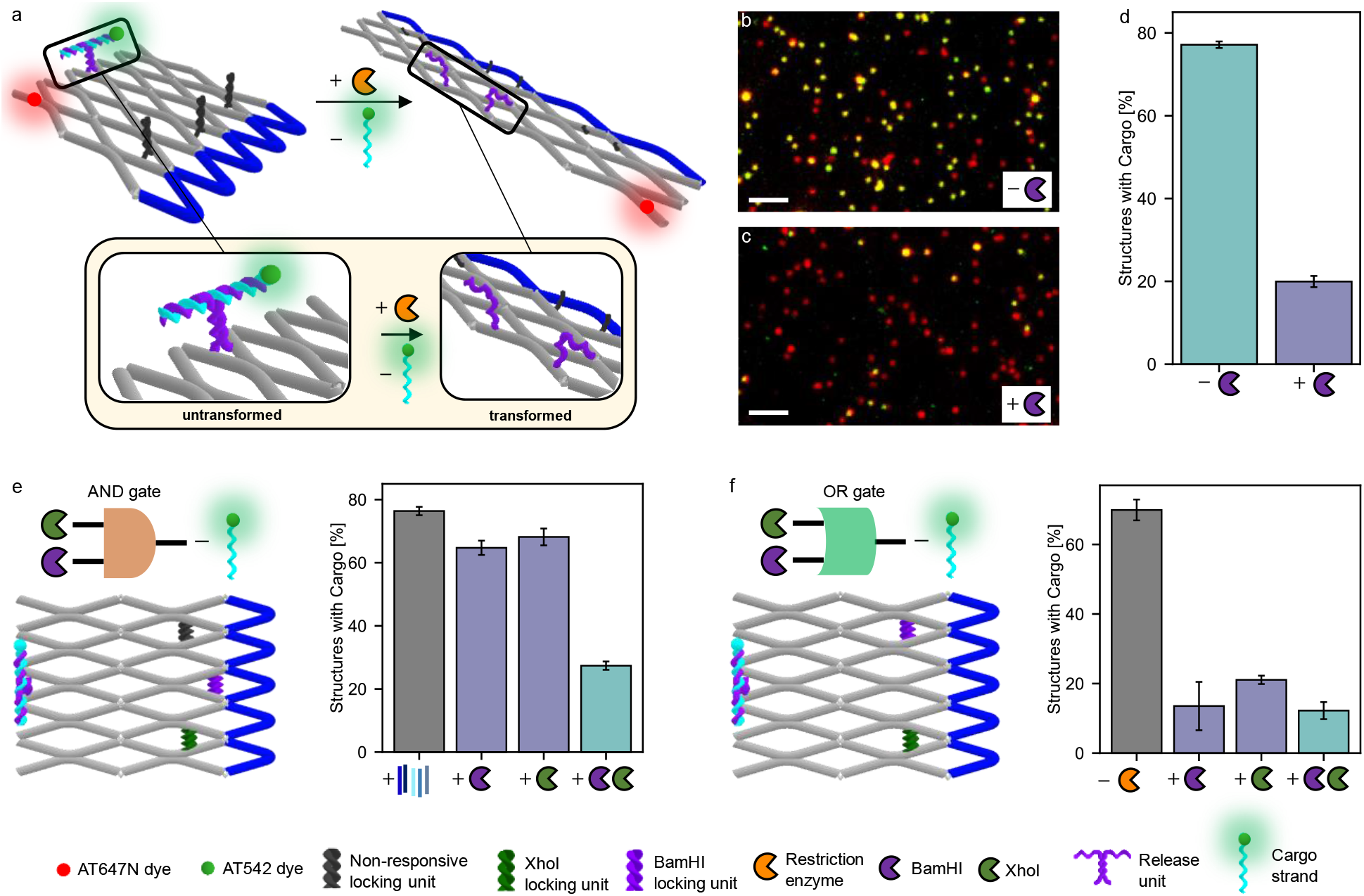
Proximity-induced output operation generated upon molecular inputs by DNA origami array nanorobot. (a) Schematic representation of DNA origami array structures which release a cargo DNA strand in response to the activity of BamHI. (b,c) TIRF images of the DNA origami array nanodevice after incubation (b) without and (c) with BamHI. (d) Corresponding fractions of DNA origami array nanodevices with a bound cargo DNA strand. (e,f) Schematic representation of DNA origami array nanodevices computing the controlled release of a cargo in response to the presence of combinations of XhoI and BamHI and corresponding fractions of DNA origami array nanodevices with a bound cargo DNA strand for an AND (e) and an OR (f) logic gate. In addition to possible inputs, all DNA origami arrays are incubated with fuel DNA strands 1-5. Error bars represent the standard deviation of the fractions calculated from three TIRF images. Scalebar: 3 μm.

First, we combine the cargo release unit with restriction enzyme locking units specific to the activity of BamHI. Here, the fraction of structures carrying a DNA cargo strand is determined from the colocalization yield with an ATTO647N dye additionally incorporated in the DNA origami.

After assembly, DNA origami arrays near quantitatively carry the cargo DNA strand (Supplementary Fig. 13). Upon incubation with and without enzyme, the fraction of structures carrying the cargo drops from 98% to 75% and 20%, respectively (Fig. 4b-d). While this indicates some unspecific cargo release, the majority of all cargo is only released in presence of the restriction enzyme. We attribute the unspecific loss of the cargo strand to the decreased stability at the incubation temperature of 37 °C (Supplementary Note 2, Supplementary Fig. 13). As we achieved both the implementation of logic gates and variable input-output combinations, we then use the modularity of our system to combine all of the above. We design structures which compute the controlled release of cargo in response to combinations of XhoI and BamHI activity. In both AND and OR logic configuration, a high fraction of cargo is released only when the gate gives a positive response (Fig. 4e,f, Supplementary Fig. 14, 15).

### Constructing networks of multiple inputs and outputs

Analogously to the input units, SEPP also enables incorporating multiple output units. This can be used to amplify these outputs as we demonstrate exemplarily for the fluorescence onset unit (Supplementary Fig. 16) or to perform different events consecutively (Supplementary Fig. 17). Here, we use timing units based on a DNA lock to control order and timing between different events.^21,22^ Timing units slightly heighten the activation barrier at a certain intermediate step in the transformation cascade, and thus introduce a time lag between operations placed on antijunctions transforming before and after them. We demonstrate this exemplarily by positioning a timing unit between two fluorescence onset units. In absence of the timing unit, the fluorescence onset units light up simultaneously upon activation by restriction enzyme activity. If a timing unit is incorporated, a time lag between both fluorescence onsets is achieved for the majority of all structures. The order in which the fluorescence onset units light up can be reversed by switching the antijunction nodes they are incorporated in (Supplementary Fig. 17).

We then set out to make full use of our designed software framework and combine all developed unit types in a single system, using ssDNA and restriction enzyme input units, fluorescence on- and offset output units (Supplementary Fig. 18) as well as a timing unit (Fig. 5a). In the absence of inputs, SEPP shows only a red fluorescence signal (Fig. 5b,c). Upon activation by a ssDNA input and XhoI restriction enzyme activity combined to a logic AND gate, SEPP switches off the red fluorescent signal (Fig. 5b,c). Depending on whether an additional timing unit is incorporated or not, a green fluorescent signal lights up either after the red fluorescence offset occurs or simultaneous to it (Fig. 5b-g). This good agreement between programmed function and execution exemplifies the control achievable in programmed systems using SEPP.

**Figure 5.**
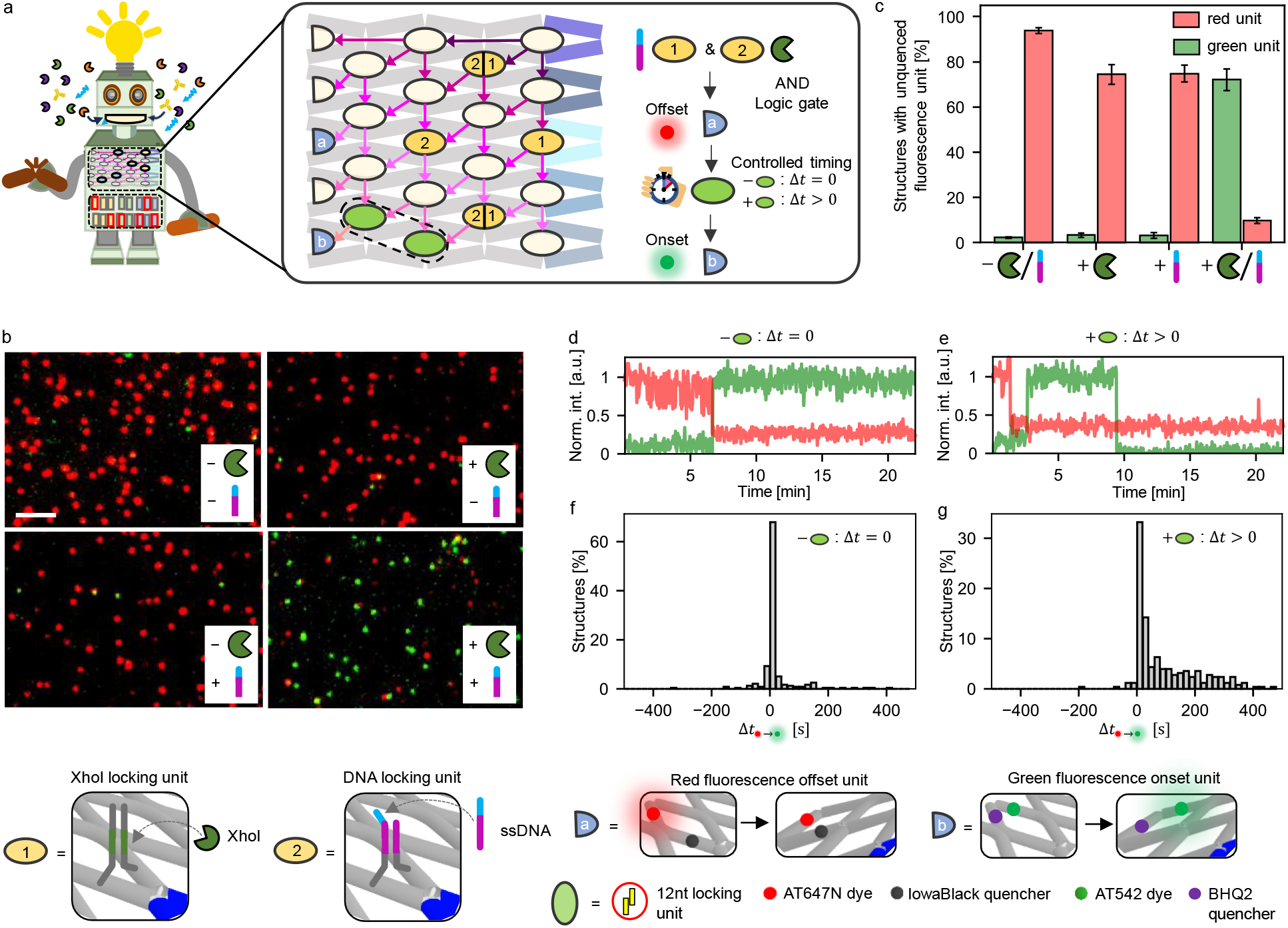
DNA origami array nanorobot performing output operations under temporal control after processing a Boolean logic AND gate based on restriction enzyme activity and ssDNA binding. (a) Schematic design of a DNA origami array nanorobot programmed to perform a series of operations with the corresponding plan of action. (b) TIRF images of the DNA origami array nanorobot after incubation with different combinations of restriction enzyme and/or ssDNA (c) Corresponding fraction of structures with unquenched green and red fluorescence units. Error bars represent the standard deviation in the fractions calculated from three TIRF images. (d,e) Exemplary single-molecule transients for structures without (d) and with (e) a timing unit incorporated. (f,g) Time between the occurrence of the red fluorescence offset and the green fluorescence onset for structures without (f) and with (g) a timing unit incorporated. In addition to possible inputs, all DNA origami arrays are incubated with fuel DNA strands 1-5. Scalebar: 3 μm.

## Conclusion

We demonstrate the rational development of DNA origami nanorobots using the network of coupled two-state systems offered by DNA origami arrays as a programmable hardware and fuel DNA strands as energy source which spring-load the system with varying degrees of tension. By designing a software package which defines different units, we show how a wide range of functionalities can be encoded into any of the network’s two-state systems and coupled together through the network’s energy landscape. We design input units responsive to ssDNA, light, restriction enzyme activity and antibodies that either activate and/ or inhibit the DNA origami array transformation. By combining multiple units on a single DNA array structure, we develop strategies to tune the responsive input concentration window, shifting K_1/2_ over 3000-fold while evolving cooperativity. We use the same strategy to incorporate Boolean logic gating, creating specific responses for different input combinations, both with inputs of the same and different molecular classes. Subsequently, we demonstrate the potential of the modularity of this software-hardware combination. We program a nanorobot that combines activation by different inputs with Boolean logic gating and a sequence of output operations such as (amplified) fluorescence output or cargo release conducted in a predefined order and under temporal control and demonstrate its proper functionality.

Overall, we expect that expanding our hardware by using different DNA origami arrays and software packages by including further proximity-based operations will be straight-forward. Including further input and output software units that – besides the demonstrated cargo release and fluorescence on-/offset – enable e.g. the on-demand onset of catalysis or cargo uptake,_19_ will pave the way for a broad range of applications in the fields of clinical diagnosis and – by implementation of arising DNA origami stabilization strategies_23–25_ – also therapeutics.

## Resource availability

### Lead contact

Philip Tinnefeld: Department of Chemistry and Center for NanoScience, Ludwig-Maximilians-Universität München, Butenandtstr. 5-13, 81377 München, Germany.

## Materials and Methods

### Synthesis of DNA origami arrays

DNA origami structures are designed using the open-source software caDNAno2 _26_ and assembled and purified using published protocols _27_ For the exact sequences of all unmodified and modified DNA staple strands used to fold the DNA origami structures see Supplementary Tables 1-10. DNA staple strands are purchased from Eurofins Genomics GmbH (Germany) and Integrated DNA Technologies (USA).

For DNA origami folding, 10 nM of in house produced p1800 scaffold (Supplementary Note 3) in 1xTAE (400 mM Tris, 400 mM acetic acid, 10 mM EDTA, pH 8) containing 12.5 mM is mixed with a 10-fold excess of all unmodified and a 30-fold excess of all modified oligonucleotides, respectively. The mixture is heated to 65 °C in a thermocycler and kept at this temperature for 15 min before being cooled down to 25 °C with a temperature gradient of – 1 °C min_-1_. Folded DNA origamis are purified from excessive staple strands by gel electrophoresis. All gels are ran using a 1.5% agarose gel, 1xTAE containing 12.5 mM MgCl_2_ for 2 hours at 6 V/cm. The target band containing DNA origami is cut from the gel and DNA origami solution extracted from the band via squeezing.

### Sample preparation on the coverslip for single-molecule widefield measurements

For chamber preparation, adhesive SecureSeal™ Hybridization Chambers (2.6 mm depth, Grace Bio-Labs, USA) are glued on microscope coverslips of 24 mm × 60 mm size and 170 μm thickness (Carl Roth GmbH, Germany). The created wells are incubated with 1 M KOH for 1 h and washed three times with 1×PBS buffer. After surface passivation by incubation with BSA-Biotin (0.5 mg/mL, Sigma Aldrich, USA) for 10 min, the surface is washed with 200 μL 1× PBS buffer. 150 μL neutravidin (0.25 mg/mL, Thermo Fisher, USA) is incubated for 10 min and then washed three times with 150 μL 1× PBS buffer. Surface immobilization is achieved via biotin-neutravidin interactions. For this, we incorporate one biotinylated DNA staple strand in the loop of the DNA origami structure during folding. The DNA origami solution is diluted with 1× TE buffer containing 750 mM NaCl to a concentration of ∼10 pM and then immobilized on the biotin-neutravidin surface via biotin-neutravidin interactions. For this, 150 μ of the DNA origami sample solution is added and incubated for 5 min. Residual unbound DNA origami is removed by washing the chambers with 150 μL 1× TE buffer containing 750 mM NaCl. The density of DNA origami on the surface suitable for single-molecule measurements is checked on a TIRF microscope. For acquisition of single-molecule fluorescence movies, an oxidizing and reducing buffer system (1x TAE, 12.5 mM MgCl_2_, 2 mM Trolox/Troloxquinone)^28^ is used in combination with an oxygen scavenging system (12 mM protocatechuic acid, 56 μM protocatechuate 3,4-dioxygenase from pseudomonas sp., 1% glycerol, 1 mM KCl, 2 mM Tris HCl, 20 μM EDTA-Na_2_*2H_2_O) to suppress blinking and photobleaching. If not stated otherwise, for acquisition of single TIRF images, no oxidizing and reducing buffer system was added.

### Reconfiguration of DNA origami array structures in response to different molecular inputs

#### ssDNA detection assay

For the detection of ssDNA, DNA origami array structures are folded with ssDNA locking units. After surface-immobilization, 50 nM fuel DNA strands in 1xTE buffer containing 750 mM NaCl and if not stated otherwise 2 uM ssDNA input are added. The samples are incubated at 37 °C for 15 min and dual-color TIRF images recorded. For titration curve measurements, samples are incubated overnight to ensure equilibrium conditions.

### Restriction enzyme activity assay

For the detection of restriction enzyme activity, DNA origami array structures are folded with restriction enzyme locking units. 50 nM of all five fuel DNA strands in 1xTAE buffer containing 12.5 mM MgCl_2_ and 2 μL of XhoI (20,000 units/ml, New England BioLabs, USA), StuI (10,000 units/ml, New England BioLabs, USA) or BamHI (100,000 units/ml, New England BioLabs, USA) are added to surface-immobilized DNA origami samples. To determine transformation yields, the structures are incubated for 15 min at 37 °C and dual-color TIRF images recorded. To measure transformation time distributions (e.g. for the incorporation of timing units), sample chambers are sealed immediately after addition of the enzymes and the photostabilization system and dual-color movies of the DNA origami arrays acquired for 20 min at 37 °C.

### Light detection assay

For the detection of light of 365 nm, DNA origami structures are folded with a light-responsive locking unit. 50 nM of all five fuel DNA strands in 1xTAE buffer containing 12.5 mM MgCl_2_ are added to surface-immobilized DNA origami samples and the samples illuminated with light of 365 nm for 5 min. After subsequent incubation at 37 °C for 15 min, dual-color TIRF images are recorded.

### Anti-Dig antibody detection assay

For the detection of anti-Dig antibodies, DNA origami array structures are folded with two Dig recognition elements. 50 nM of fuel DNA strands 1 and 2 in 1xTE buffer containing 750 mM NaCl and 100 nM anti-Dig antibodies (Rb Monoclonal, Thermo Fisher Scientific, cat#: 700772, PRID: AB_2532342) are added to surface-immobilized DNA origami samples. Samples are incubated at 37 °C for 15 min and dual-color TIRF images recorded.

### Boolean logic gating and cargo release assays

For the measurement of Boolean logic gates and cargo release, surface-immobilized DNA origami structures are incubated with 50 nM fuel DNA strands 1-5 in 1xTAE buffer containing 12.5 mM MgCl_2_ and the different restriction enzyme/ light inputs at 37 °C as described above and dual-color TIRF images recorded.

### Nanorobot measurement combining multiple inputs with multiple outputs

For the nanorobot measurement which combines multiple inputs with multiple outputs (Fig. 5), surface-immobilized DNA origami array structures are incubated with 50 nM fuel DNA strands 1-5 in 1xTAE buffer containing 12.5 mM MgCl_2_ and different combinations of 2 uL BamHI/ 2 uM ssDNA at 37 °C as described above. For determining the fraction of structures with unquenched fluorescent units, dual-color TIRF images are recorded after 15 min incubation time. For determining the time delay between the red fluorescence offset and the green fluorescence onset, sample chambers are sealed immediately after addition of both inputs and the photostabilization system and dual-color movies of the DNA origami arrays acquired for 20 min at 37 °C.

### Wide-field measurements

For detection of single-molecule fluorescence, a commercial wide-field/TIRF microscope Nanoimager from Oxford Nanoimaging Ltd. is used. Red excitation at 638 nm is realized with a 1100 mW laser, green excitation at 532 nm with a 1000 mW laser, respectively. The relative laser intensities are set to 9% for green and to 18% for red excitation. The microscope is set to TIRF illumination. Measurements are carried out at 37 °C. For quantifying transformation yields and the percentage of structures carrying cargo, dual-color fluorescence images are acquired. For recording fluorescence movies, the lasers are activated and a frame of 100 ms is taken every second separately for both excitation lasers (with a time lag of 0.5 s between them) over a measurement period of 20 min.

## Data analysis

We quantify the percentage of transformed structures by dividing the number of green (ATTO 542) and red (ATTO647N) co-localized spots by the total number of green spots from dual-color TIRF images. To account for a labelling efficiency < 100%, the percentage of co-localized spots is normalized by the percentage of co-localized spots of a DNA origami array folded with all five fuel DNA strands to calculate transformation yields. In the normalization sample, AFM imaging confirmed the full transformation of all structures in previous work.^21^ For structures with a cargo release unit incorporated, the percentage of structures carrying cargo is determined analogously.

Data processing and analysis of time-lapse movies is realized using custom-written Python scripts. Briefly, the acquired movies are first drift corrected using DNA origami structures carrying fluorophores which are in their fluorescent state throughout the whole measurement as fiducial markers. Spots appearing during the measurement are detected from the drift-corrected movies and dual-color background-subtracted fluorescence intensity transients of those spots extracted. To determine transformation times of single structures, the corresponding transients are fitted using a Hidden Markov model (HMM). Two levels corresponding to the quenched and unquenched state of the corresponding fluorescence unit are defined. For fluorescence onset (offset) units, transformation times are defined as the time a structure switches from its quenched (unquenched) state to its unquenched (quenched) state and subsequently remains in it for at least 10 s for the first time. To determine transformation time distributions, only structures showing an intensity change in both colors are considered.

For titration curve measurements, the transformation yields obtained upon incubation with different concentrations of ssDNA input [*DNA*] are calculated from dual-color TIRF images as described above. *K*_*1/2*_ and the Hill coefficient *n*_*H*_ are subsequently determined by fitting the calculated transformation yields *Y* to the modified Hill equation.

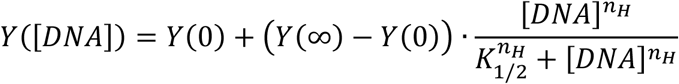

where *Y*(0) and *Y* (∞) give the start and the end points of the titration.

For determining the fraction of structures with unquenched fluorescence unit in Fig. 5, we quantify the total number of structures in a TIRF image by counting the number of both weak (quenched) and bright (unquenched) red fluorescent spots. The fraction of red unquenched fluorescence units is then calculated as the fraction of the bright red spots of the total number of red spots. As green quenched fluorescence units are not visible in TIRF images, the fraction of green unquenched fluorescence units is calculated by dividing the number of green spots by the total number of red spots and subsequently normalizing them by the fraction of colocalized red and green spots in fully transformed DNA origami arrays.

## Supporting information

Suppplementary Information

## Acknowledgments

P.T. gratefully acknowledges financial support from the Federal Ministry of Education and Research (BMBF) in the framework of the Cluster4Future program (Cluster for Nucleic Acid Therapeutics Munich, CNATM) (Project ID: 03ZU1201AA) as well as the Free State of Bavaria under the Excellence Strategy of the Federal Government and the Länder through the ONE MUNICH Project Munich Multiscale Biofabrication. Y.K. acknowledges the support by the Department of Energy grant DESC0020996, the National Science Foundation grants CCF-2227399 and ECCS-2328217, and the National Institute of Health grants 1R35GM153472 and 1RM1GM145394. M.P. acknowledges the support by Studienstiftung des Deutschen Volkes. The authors thank Luna, Charly and Xaverl for their support with the measurements and Angelika, Frau Steger and Herr Ehrl for lab upkeep.

## Author contributions

All authors conceived and developed the concept. M.P., F.C. and D.W. prepared samples, performed and analyzed the measurements. Y.K. and P.T. supervised the project. All authors have written, read and approved the final manuscript.

## Data and code availability

All data and codes used in this study are available upon reasonable request from the corresponding authors.

