## Supplementary material for "Spring-loaded DNA origami arrays as energy-supplied hardware for modular nanorobots": Suppplementary Information

a

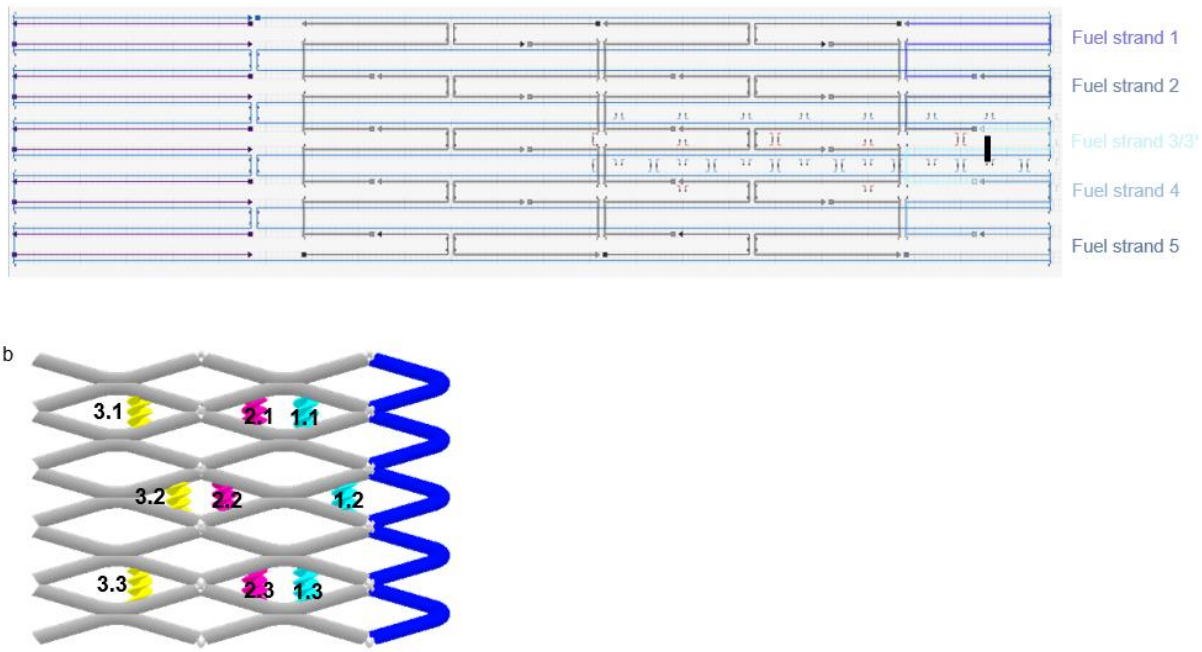


**Supplementary Figure 1. Design of the 5 × 2.5 reconfigurable DNA origami array structure in its untransformed conformation.** (a) Blue, gray, purple and blue lines represent the scaffold strand, core DNA staple strands of the structure, loop staple strands and fuel DNA strands, respectively. The DNA loop spans the structure from one end to the other but does not participate in the transformation process. By labeling one of the staple strands with biotin, we use it as an anchor point for surface immobilization via biotin-neutravidin interactions. Fuel DNA strands 1-4 all have the same length of 65 base pairs. Fuel DNA strand 5 is shorter, consisting of only 39 base pairs. Fuel strand 3* is shortened version of fuel strand 3 consisting of only 25 base pairs. The position of its 5-prime end is marked with a black line. As the transformation process is starts either at the upper right or lower right corner, this asymmetry induces a preferential transformation starting point. The longer length of fuel DNA strand 1 compared to fuel DNA strand 5 results in the transformation preferentially starting from the upper right corner. (b) Sketch illustrating the positions of the activation locking units specific to ssDNA, restriction enzyme activity and light at different anti-junctions of the DNA origami array structure.

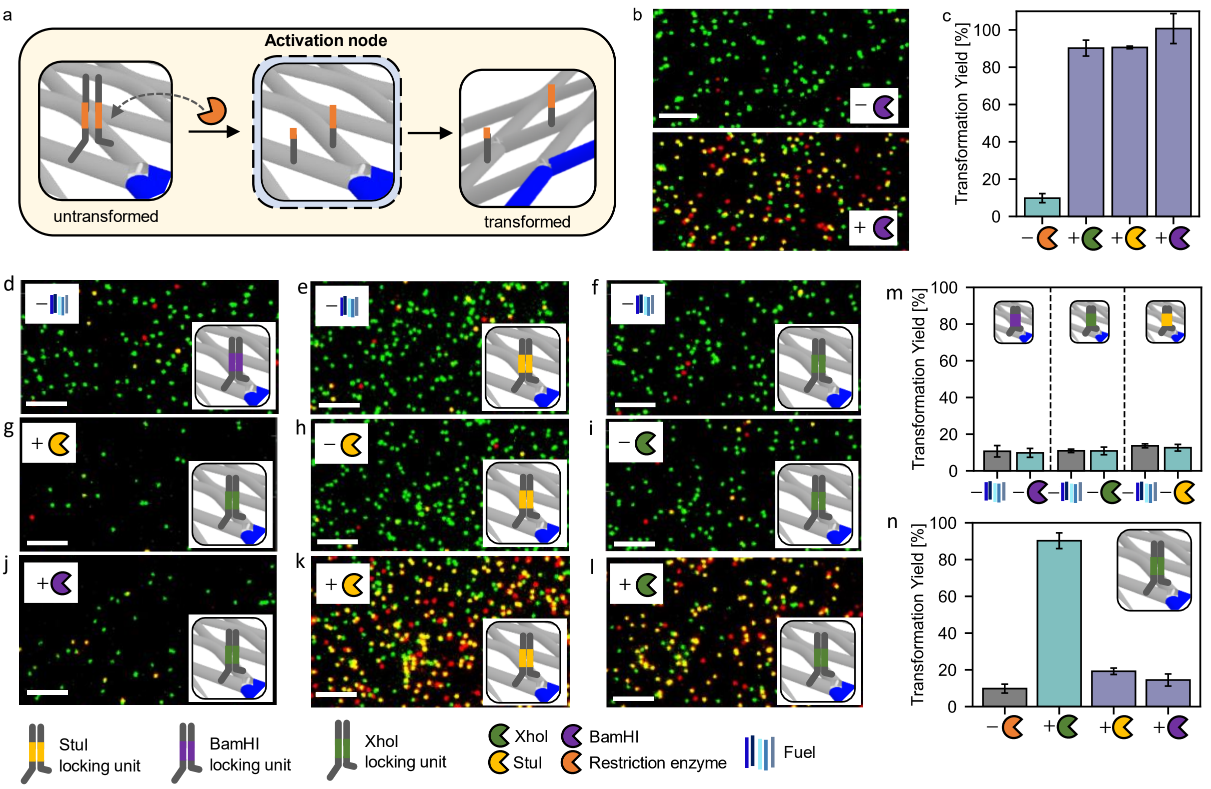


**Supplementary Figure 2. Restriction enzyme activity as input.** (a) Sketch of nodes responsive to restriction enzyme activity. Their design is based on a dsDNA lock containing the restriction enzyme-specific cleavage and binding site. The lock is cleaved in presence of active restriction enzyme. (b) Exemplary TIRF images of DNA origami array structures with three restriction enzyme locking units (positions 1.1, 1.2 and 1.3) containing the binding and cleavage site for BamHI before and after incubation with DNA fuel strands and after incubation with fuel strands and BamHI. (c) Transformation yields of DNA origami array structures with three restriction enzyme locking units containing the binding and cleavage site for BamHI, StuI and XhoI, respectively, before and after incubation with DNA fuel strands and BamHI, StuI and XhoI, respectively. (d-l) Exemplary TIRF images of DNA origami array structures with three restriction enzyme locking units before and after incubation with DNA fuel strands and after incubation with fuel strands and restriction enzymes. (m,n) Corresponding transformation yields. (m) When adding the fuel DNA strands (-Enzyme) no significant increase in transformation yield is observed, indicating that by introduction of the restriction enzyme locking units, the transformation process cannot be induced by the fuel DNA strands alone. (n) Only in presence of XhoI, DNA origami array structures with three XhoI locking units show a high transformation yield, indicating good specificity. In addition to possible inputs, all DNA origami arrays are incubated with fuel DNA strands 1-5. Error bars represent the standard deviation in the transformation yields calculated from three TIRF images. Scalebar: 4 µm.

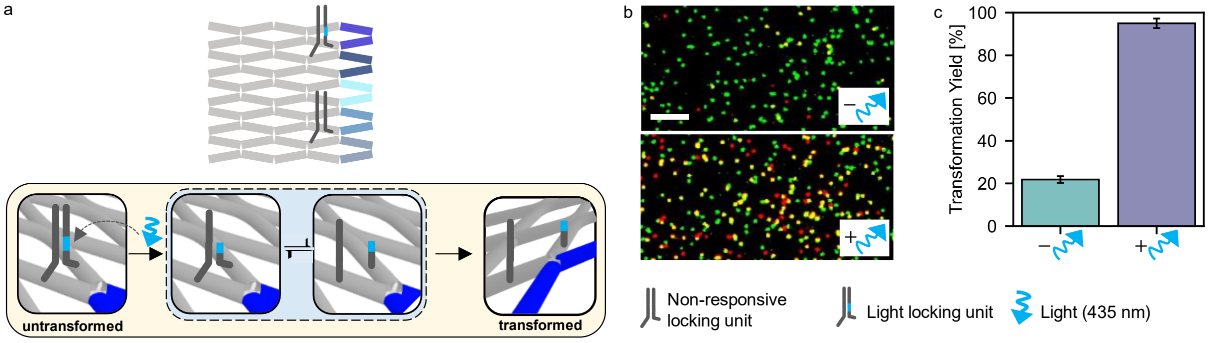


**Supplementary Figure 3. Light as input.** (a) Sketch of nodes responsive to light. The design is based on a dsDNA lock containing a light-cleavable linker which is cleaved in presence of light). (b) Exemplary TIRF images of DNA origami array structures with a light-cleavable locking unit (position 1.1) and an additional stabilization unit which stabilizes the untransformed state of the array and does not interact with any inputs (position 1.3) incubated with fuel DNA strands 1-5 in the presence and absence of light. (c) Corresponding transformation yields. Error bars represent the standard deviation in the transformation yields calculated from three TIRF images. Scalebar: 4 µm.

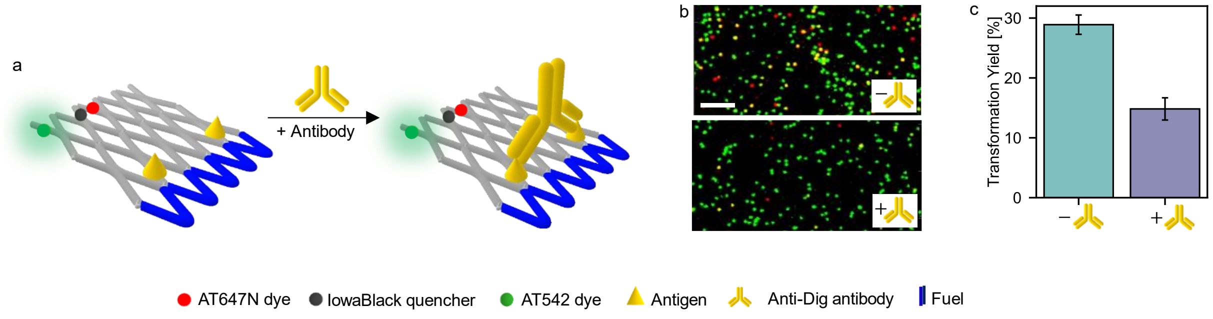


**Supplementary Figure 4. Antibody as input.** (a) Sketch of DNA origami array structure bearing two antigens as binding elements for an antibody. Bivalent binding of an antibody inhibits the transformation process. (b) Exemplary TIRF images of DNA origami array structures with Dig antigen input units before (upper) and after (middle) incubation with fuel DNA strands 1,2 and after incubation with anti-Dig antibodies and fuel DNA strands 1,2 (lower). (b) Corresponding transformation yields. Error bars represent the standard deviation in the transformation yields calculated from three TIRF images. Scalebar: 4 µm.

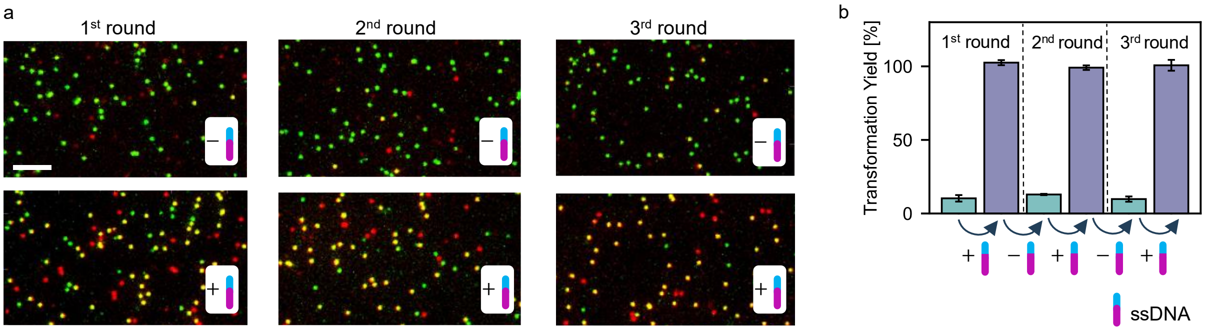


**Supplementary Figure 5. Reversibility of the transformation process activated by ssDNA input.** (a) Exemplary TIRF images of DNA origami array structures with six ssDNA activation units (positions 1.1-1.3, 3.1-3.3) incorporated throughout three rounds of incubation with ssDNA input, washing and incubation without ssDNA inputs. The measurements are carried out in the presence of fuel DNA strands 1,2,4. (b) Corresponding transformation yields. In addition to ssDNA as a possible input, the DNA origami arrays are incubated with fuel DNA strands 1-5 throughout all steps. Error bars represent the standard deviation in the transformation yields calculated from three TIRF images. Scalebar: 4 µm.

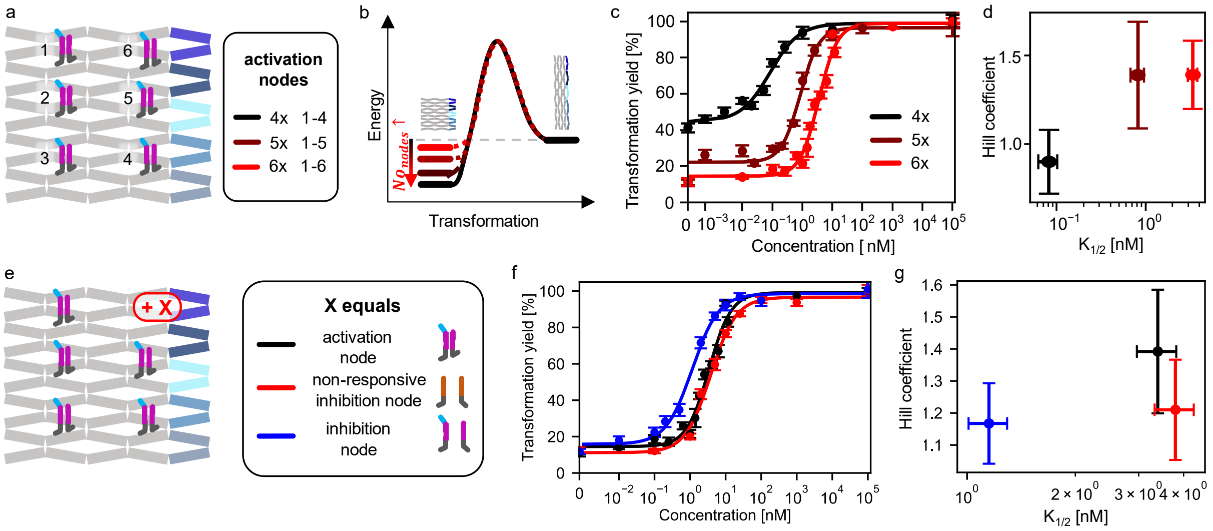


**Supplementary Figure 6. Further parameters to tune the responsive concentration window of the ssDNA input.** (a-d) Effect of the number of incorporated ssDNA activation units. (a) Design of the DNA origami array carrying different numbers of ssDNA activation units. (b) When increasing the number of ssDNA activation units, the untransformed state of the DNA origami system is stabilized, decreasing the tension in the spring-loaded system. (c-d) This shifts the responsive window to higher concentrations while simultaneously increasing the Hill coefficient. (c) (e-g) Effect of the additional incorporation of different units to a DNA array carrying five ssDNA activation units. (e) Design of the DNA origami array carrying five ssDNA activation units and additionally a variable unit X. Activation and inhibition units stabilize the untransformed and the transformed conformation of the antijunction node they are placed on, respectively. Their stabilizing effect is removed by binding of the ssDNA input to the unit. In contrast, non-responsive inhibition units stabilize the transformed conformation of the antijunction node they are placed on but do not interact with the ssDNA input. (f) Varying the unit X shifts the responsive window. (g) As expected, the addition of a non-responsive inhibition unit and an inhibition unit results in lower Hill coefficients than the addition of another activation unit. Also, the introduction of these units shifted $K_{1/2}$ to different extents. In addition to ssDNA as a possible input, all DNA origami arrays are incubated with fuel DNA strands 1-5. Error bars in (c,f) represent the standard deviation in the transformation yields calculated from at least three TIRF images. Error bars in (d,g) represent the fit error to the curves fitted in (c,f).

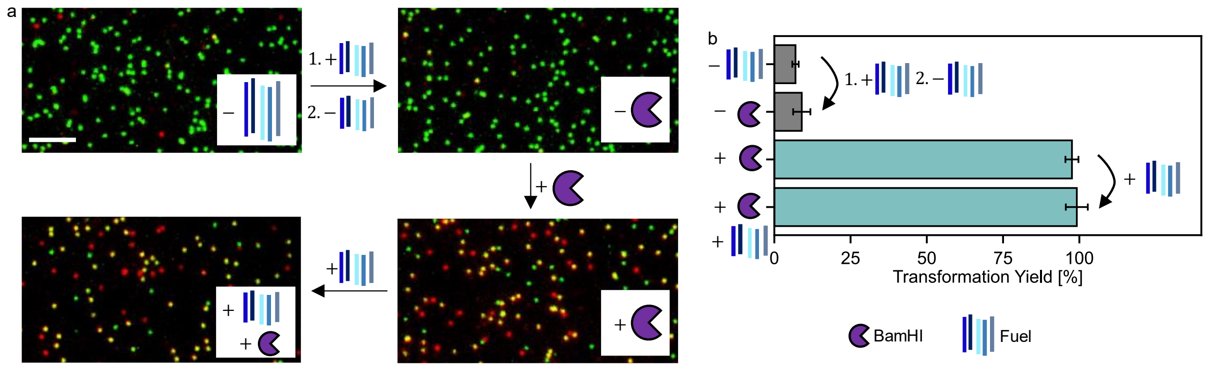


**Supplementary Figure 7. Pre-loading DNA origami array structures with fuel DNA strands.** We first incubate a DNA origami array structures with three BamHI locking units (positions 1.1, 1.2 and 1.3) with the fuel DNA strands before removing unbound fuels in solution by five washing steps. Subsequent addition of BamHI results in a near quantitative transformation of all structures, demonstrating the successful pre-loading with fuel strands and creation of the pre-tensioned state. The near quantitative transformation is confirmed by again adding the fuel DNA strands at the end of the assay which does not result in a significant increase in transformation yield. Thus, we conclude that pre-loading the fuel DNA strands and their attached energy in a quantitative manner is possible (a) Exemplary TIRF images of the DNA origami arrays before incubation with fuel DNA strands (upper left), after incubation with fuel DNA strands 1-5 which then are removed from solution (upper right), after incubation with BamHI (lower right) and again adding the fuel DNA strands 1-5 (lower left). (b) Corresponding transformation yields. Error bars represent the standard deviation in the transformation yields calculated from three TIRF images. Scalebar: 4 μm.

**Supplementary Note 1. Effect of restriction enzyme input units specific for XhoI placed at different positions as well as combinations of these units.**

For studying the inhibition effect of locking units on the overall transformation process, we place different numbers of locking units specific for XhoI at different antijunctions nodes in the system (see Fig. S8a, S8b). Depending on which antijunction the locking units are placed on, we observe different efficiencies in its inhibition of the reconfiguration process. We find that the position-dependency is directly linked to the energy landscape of the reconfiguration process: the further right the antijunction of the input unit is positioned in the DNA array, the earlier it reconfigures its conformation in the transformation process and the larger is the effect of the corresponding locking unit on the overall transformation yield. This is in good agreement with the proposed energy landscape which is tilted more strongly downwards towards the end of the transformation. Thus, in the beginning, an additional energy barrier of the same size has a larger effect than in a downhill tilt. Due to the identical design of the developed restriction enzyme locking units specific for XhoI, StuI, BamHI we expect that the obtained position-dependencies to be transferable to StuI and BamHI. We use the position-dependency to define design strategies for Boolean logic AND and OR gates (see Fig. S8c-f).

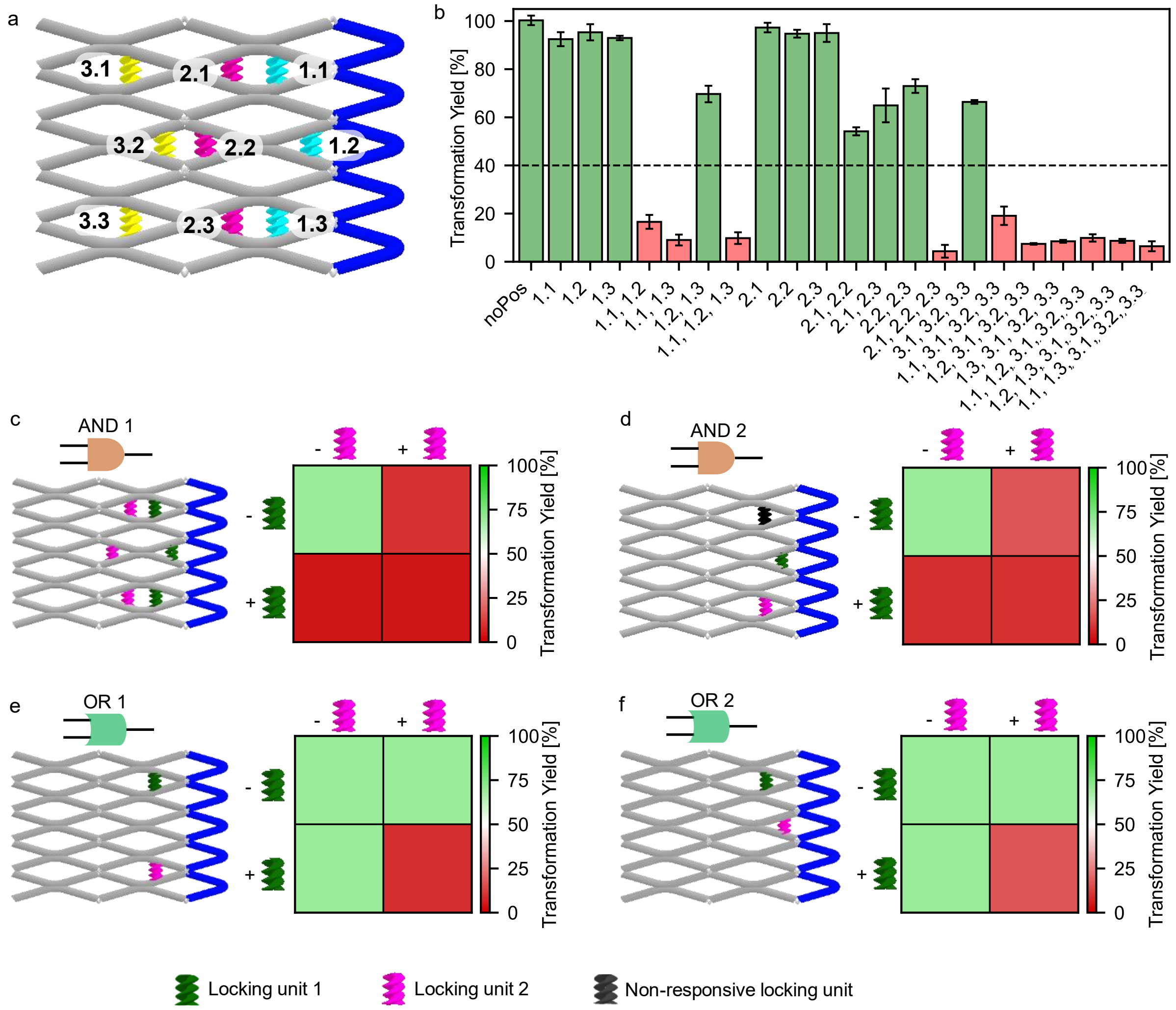


**Supplementary Figure 8. Effect of locking units specific for XhoI placed at different positions as well as combinations of these units.** (a) Sketch illustrating the nomenclature of the XhoI locking units positioned at different anti-junctions of the DNA origami array structure. (b) Transformation yields of DNA origami array structures with different combinations of XhoI locking units incorporated. The yields are obtained after incubation with the fuel DNA strands but in absence of restriction enzyme. DNA origami arrays with transformation yields above 40% (dashed line) are considered successfully transformed and marked in green. DNA origami arrays exhibiting lower transformation yields are marked in red. (c-f) Two possible DNA origami array configurations for Boolean logic AND (c,d) and OR (e,f) gates using two types of restriction enzyme as inputs. The corresponding restriction enzyme locking units are marked in the designs in pink and forest green (left sketches). Non-responsive locking units which stabilize the untransformed state of the antijunction node they are placed on and are inactive towards either of the restriction enzymes are marked in black. The transformation yields obtained for DNA origami arrays with (+) and without (-) the locking units 1/ 2 incorporated indicate the possibility of implementing Boolean logic gates based on restriction enzyme activity (right plots). Transformation yields are the same as in (b). When applying the designs to restriction enzyme activities, the absence (presence) of a locking unit corresponds to the presence (absence) of a restriction enzyme which cleaves the unit. As such, the configurations are expected to correspond to AND (c,d) gates and OR (e,f) gates with respect to the activity of restriction enzymes. The designs of (c) and (e) are used in Figure 3 of the main paper. Multi-level Boolean logic gates are designed analogously. All DNA origami arrays are incubated with fuel DNA strands 1-5. Error bars represent the standard deviation in the transformation yields calculated from three TIRF images.

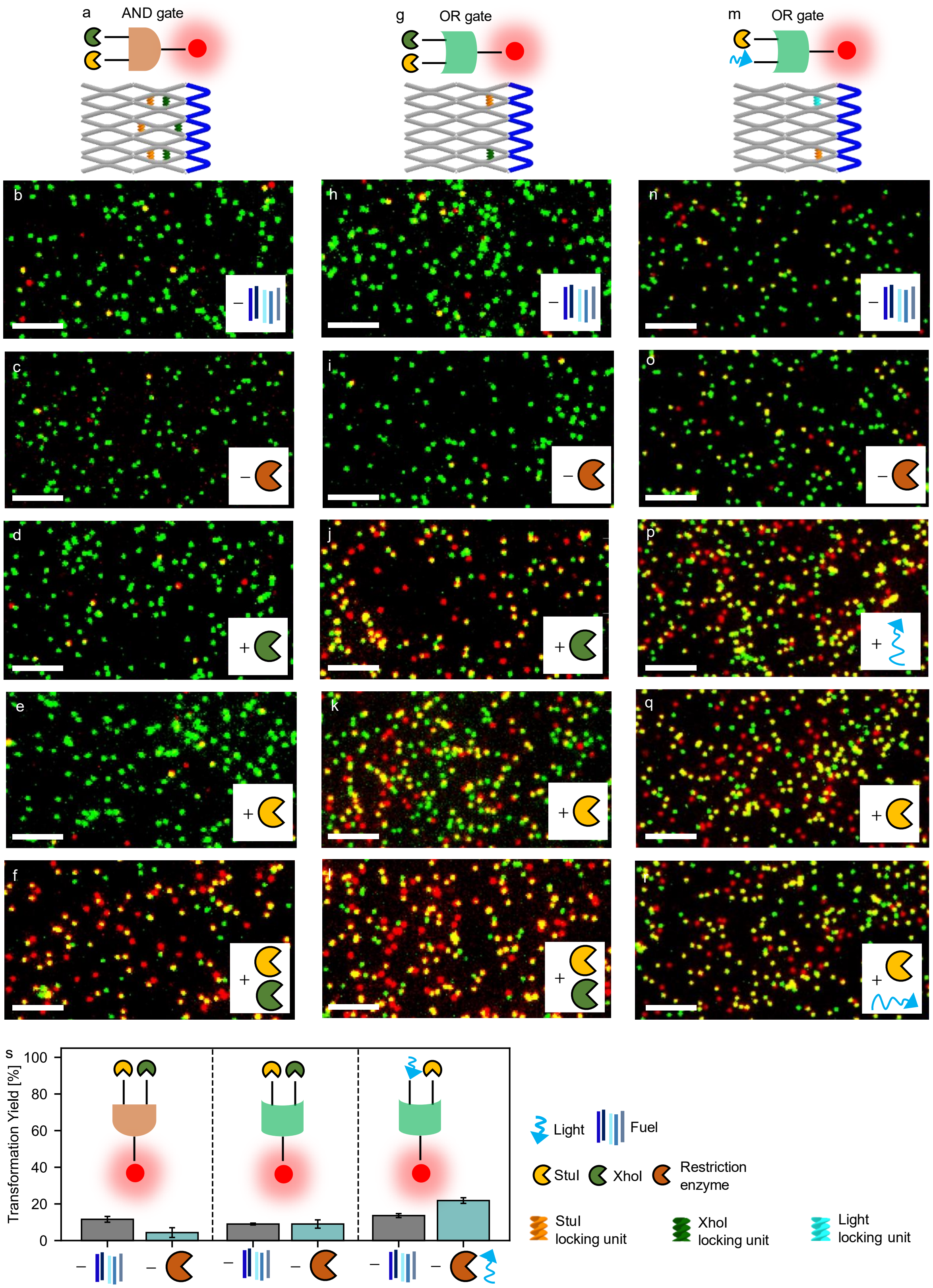


**Supplementary Figure 9. Processing one-level Boolean logic gates.** (a-f) Schematic representation of DNA origami array structures with (a) an AND logic gate responsive to combinations of StuI and XhoI and (b-f) exemplary TIRF images before and upon addition of different inputs. (g-l) Schematic representation of DNA origami array structures with (g) an OR logic gate responsive to combinations of StuI and XhoI and (h-l) exemplary TIRF images before and upon addition of different inputs. (m-q) Schematic representation of DNA origami array structures with (m) an AND logic gate responsive to combinations of StuI and light and (m-q) exemplary TIRF images before and upon addition of different inputs. (s) Corresponding transformation yields. In addition to possible inputs, all DNA origami arrays are incubated with fuel DNA strands 1-5. Error bars represent the standard deviation in the transformation yields calculated from three TIRF images. Scalebar: 4 µm.

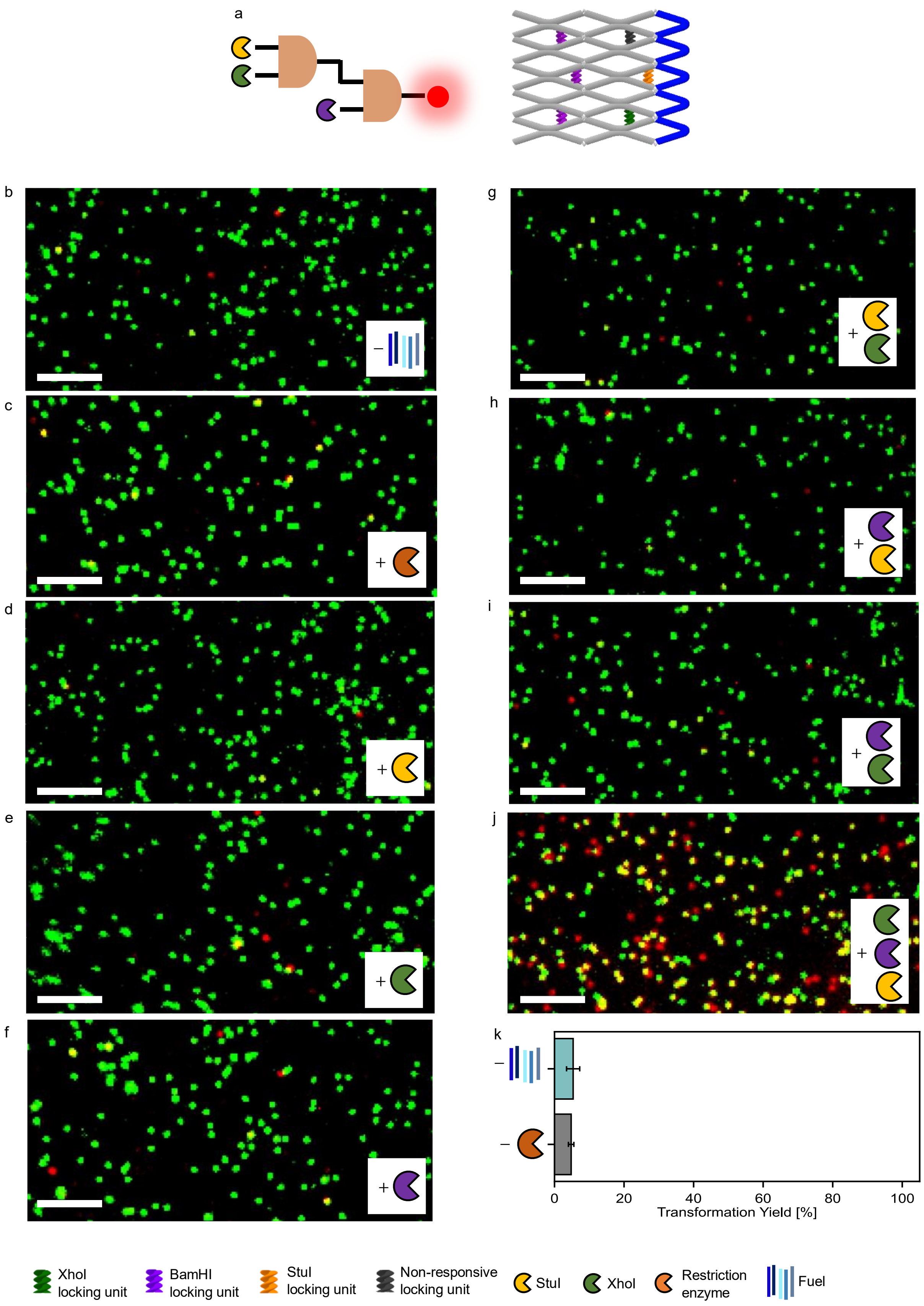


**Supplementary Figure 10. DNA origami array nanodevices processing a 3xAND gate.** (a-j) Schematic representation of DNA origami array structures with a 3xAND logic gate responsive to combinations of StuI, XhoI and BamHI and (b-j) exemplary TIRF images before and upon addition of different inputs. (k) Transformation yields obtained before and after incubation with fuel DNA strands but without restriction enzymes. In addition to possible inputs, all DNA origami arrays are incubated with fuel DNA strands 1-5. Error bars represent the standard deviation in the transformation yields calculated from three TIRF images. Scalebar: 4 μm.

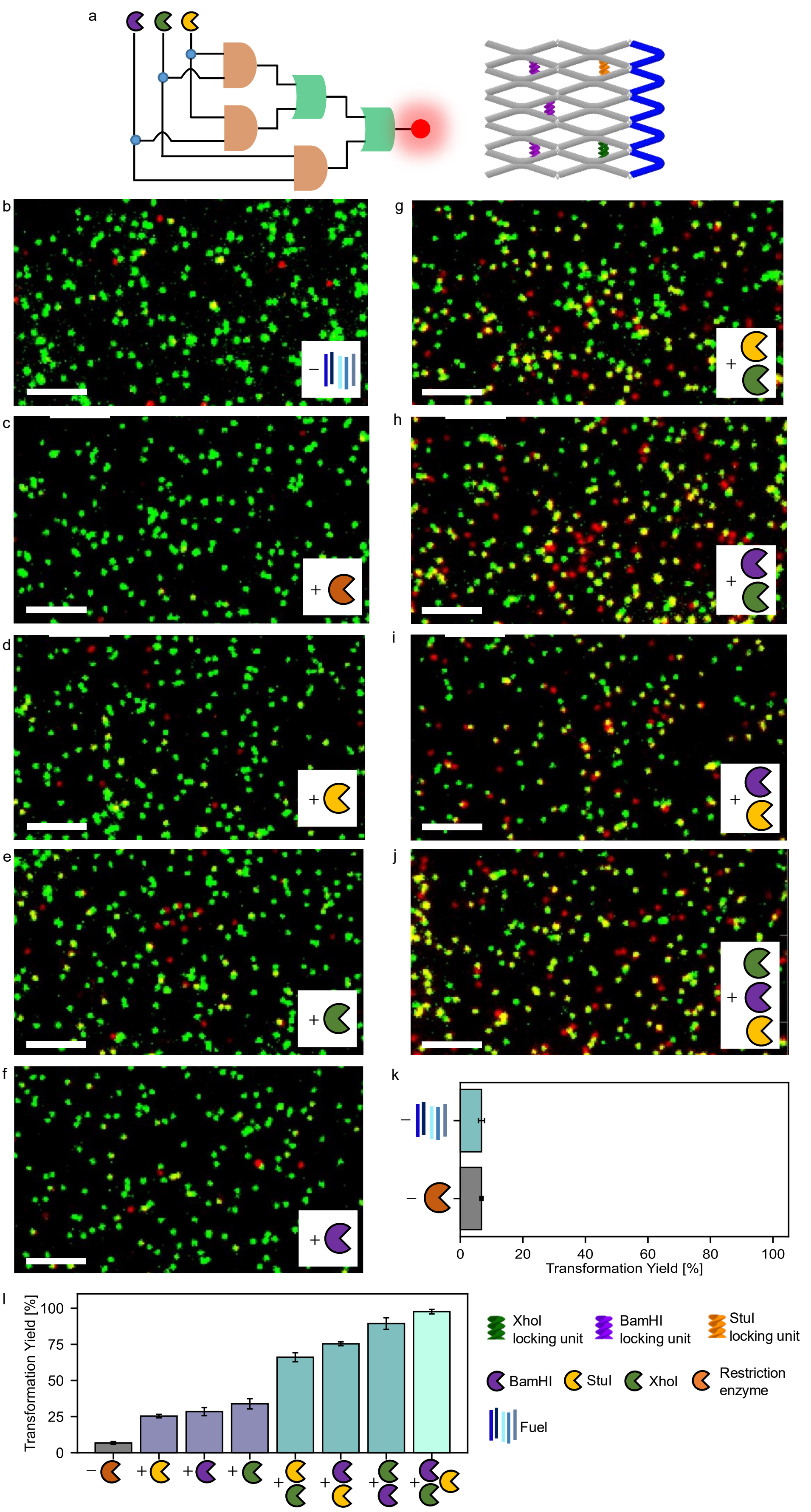


**Supplementary Figure 11. DNA origami array nanodevices processing a multi-level logic gate consisting of a series of three AND gates and two OR gates.** (a) Schematic representation of DNA origami array structures with a multi-level logic gate responsive to combinations of StuI, XhoI and BamHI and (b-j) exemplary TIRF images before and upon addition of different inputs. (k) Transformation yields obtained before and after incubation with fuel DNA strands but without restriction enzymes. (l) Transformation yields before and after incubation with fuel DNA strands and restriction enzymes. In addition to possible inputs, all DNA origami arrays are incubated with fuel DNA strands 1-5. Error bars represent the standard deviation in the transformation yields calculated from three TIRF images. Scalebar: 4 μm.

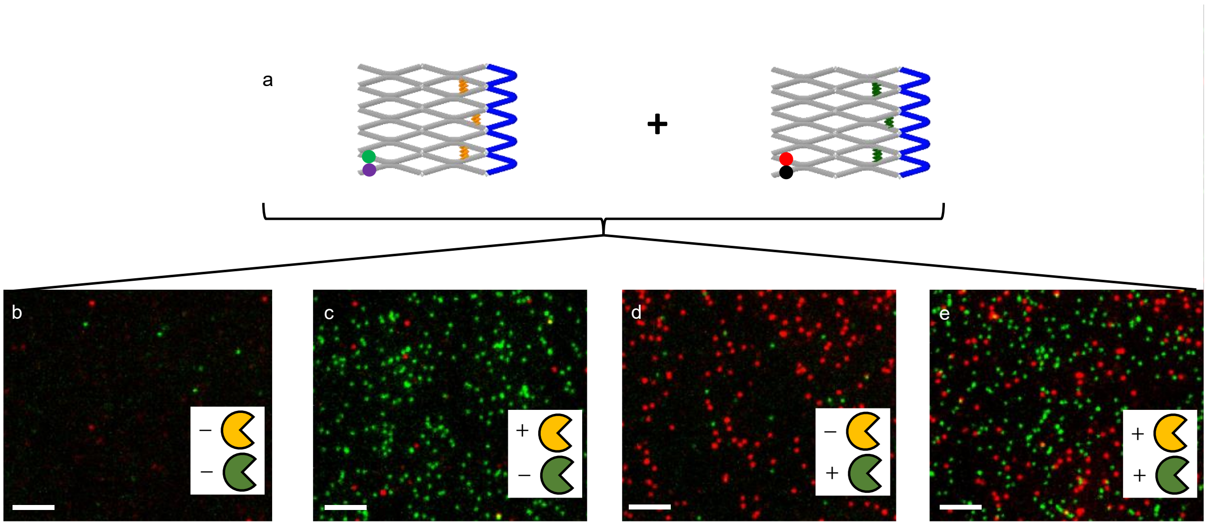


**Supplementary Figure 12. Multiplexing with DNA origami arrays.** (a) Sketch of the two different DNA origami array designs used for multiplexing. The DNA origami array responsive to StuI restriction enzyme activity (left panel) carries a green fluorescence onset unit whereas the DNA origami array responsive to XhoI restriction enzyme activity (right panel) carries a red fluorescence onset unit. By immobilizing both DNA origami arrays on the same surface, multiplexing is achieved by spectral separation. (b-e) Exemplary TIRF images of surfaces bearing both DNA origami array structures upon incubation with and without StuI (yellow) and XhoI (green) restriction enzymes. Only in the presence of the corresponding enzymes, green and/ or red spots appear. In addition to possible inputs, all DNA origami arrays are incubated with fuel DNA strands 1-5. Scalebar: 4 µm.

### Supplementary Note 2. Cargo Release Unit

The cargo release unit is formed by two ssDNA strands protruding from domains of two neighboring antijunctions. The ssDNA strands are placed on the antijunction domains such that they are in close proximity in the untransformed conformation of the DNA array and further apart in its transformed conformation. They each contain a 10 nt non-complementary linker sequence followed by a 9 nt complementary sequence which – if both strands are in close proximity – forms a stem. The stem is followed by a 10 nt non-complementary sequence on each strand to which an ATTO542-labelled ssDNA strand containing a 20-nt complementary sequence is hybridized during DNA origami array folding.

Transformation of the antijunctions carrying the cargo release unit then results in the spatial separation of the two ssDNA strands forming the cargo release unit. The stem dehybridizes resulting in a weakened affinity of the cargo strand to the release unit and subsequently to its release.

During measurements also a certain degree of unspecific cargo release is observed. Both upon incubation with and without fuel DNA strands but without restriction enzyme, we note a decrease in the fraction of structures carrying cargo. As this decrease occurs both upon incubation with and without fuel DNA strands, we do not attribute the unspecific cargo release mainly to the addition of fuel DNA strands but to the heightened incubation temperature of 37 °C (Supplementary Fig. 13)

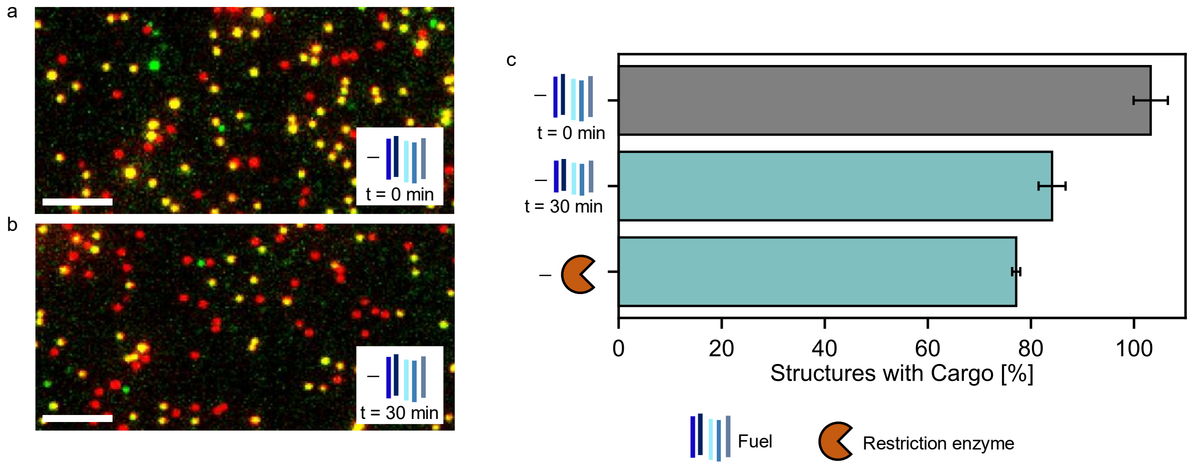


**Supplementary Figure 13. Unspecific release of a cargo DNA strand from DNA origami array structures upon incubation at 37°C and upon incubation without restriction enzyme.** (a) Exemplary TIRF images of DNA origami array structures (a) before and (b) after 30 min incubation without fuel DNA strands at 37°C. (c) Corresponding fraction of structures with cargo DNA strand before and upon incubation without and with fuel DNA strands but without restriction enzyme. After incubation, we note a decrease in the fraction of structures carrying cargo both without and with fuel DNA strands but without restriction enzyme. Thus, we do not attribute the unspecific cargo release mainly to the addition of the fuel DNA strands but to the heightened incubation temperature of 37 °C. In addition to possible inputs, all DNA origami arrays are incubated with fuel DNA strands 1-5. Error bars represent the standard deviation in the fractions calculated from three TIRF images. Scalebar: 4 µm.

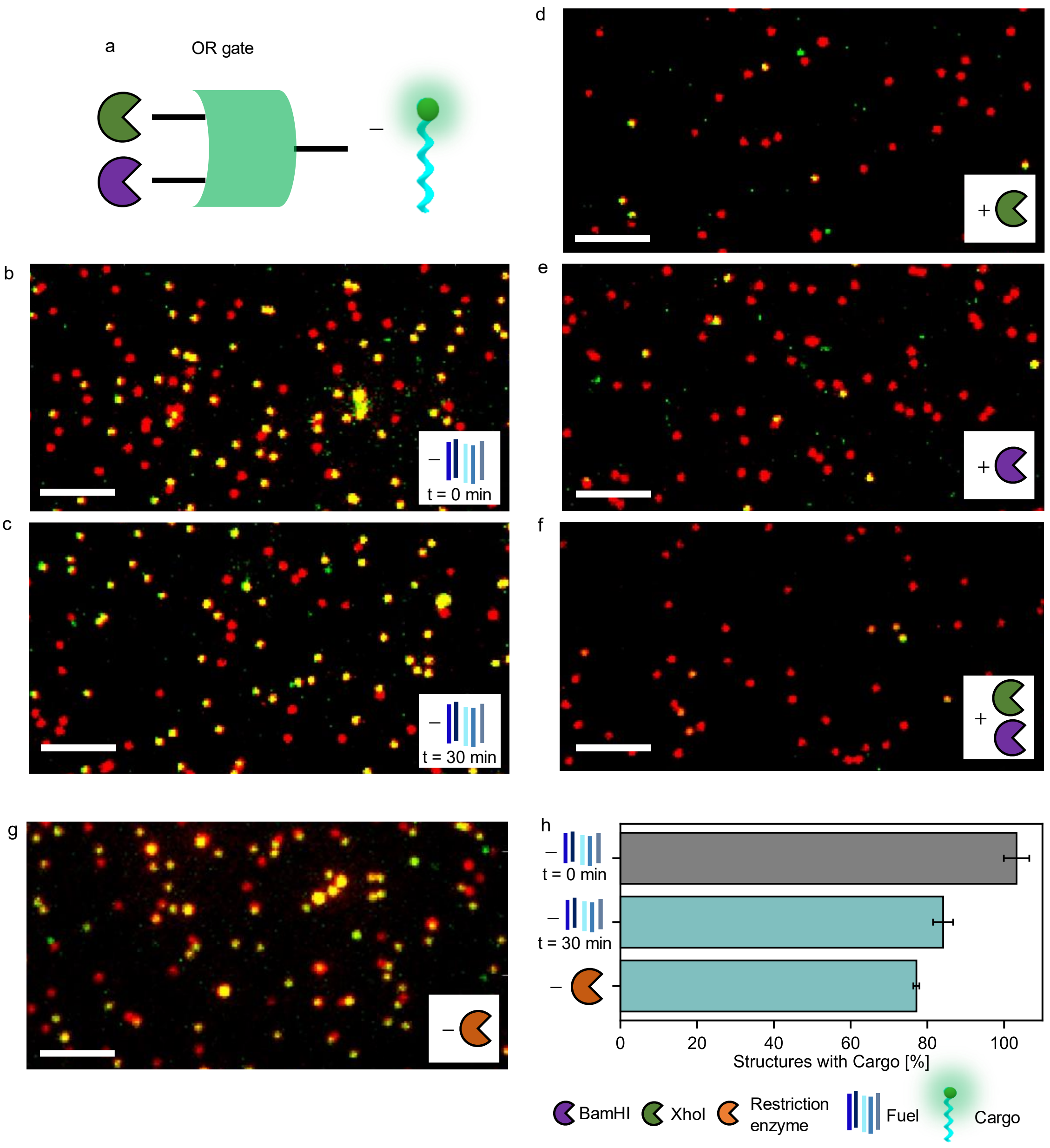


**Supplementary Figure 14. DNA origami arrays with OR logic gate releasing a cargo DNA strand in response to the combination of different restriction enzymes.** (a) Schematic representation of a OR logic gate which releases a cargo DNA strand in response to the activity of BamHI and XhoI. (b-g) Exemplary TIRF images of DNA origami array structures before and after incubation with different inputs. (h) Fraction of structures with cargo before and after incubation without and with fuel DNA strand but without restriction enzymes. In addition to possible inputs, all DNA origami arrays are incubated with fuel DNA strands 1-5. Error bars represent the standard deviation in the fractions calculated from three TIRF images. Scalebar: 4 µm.

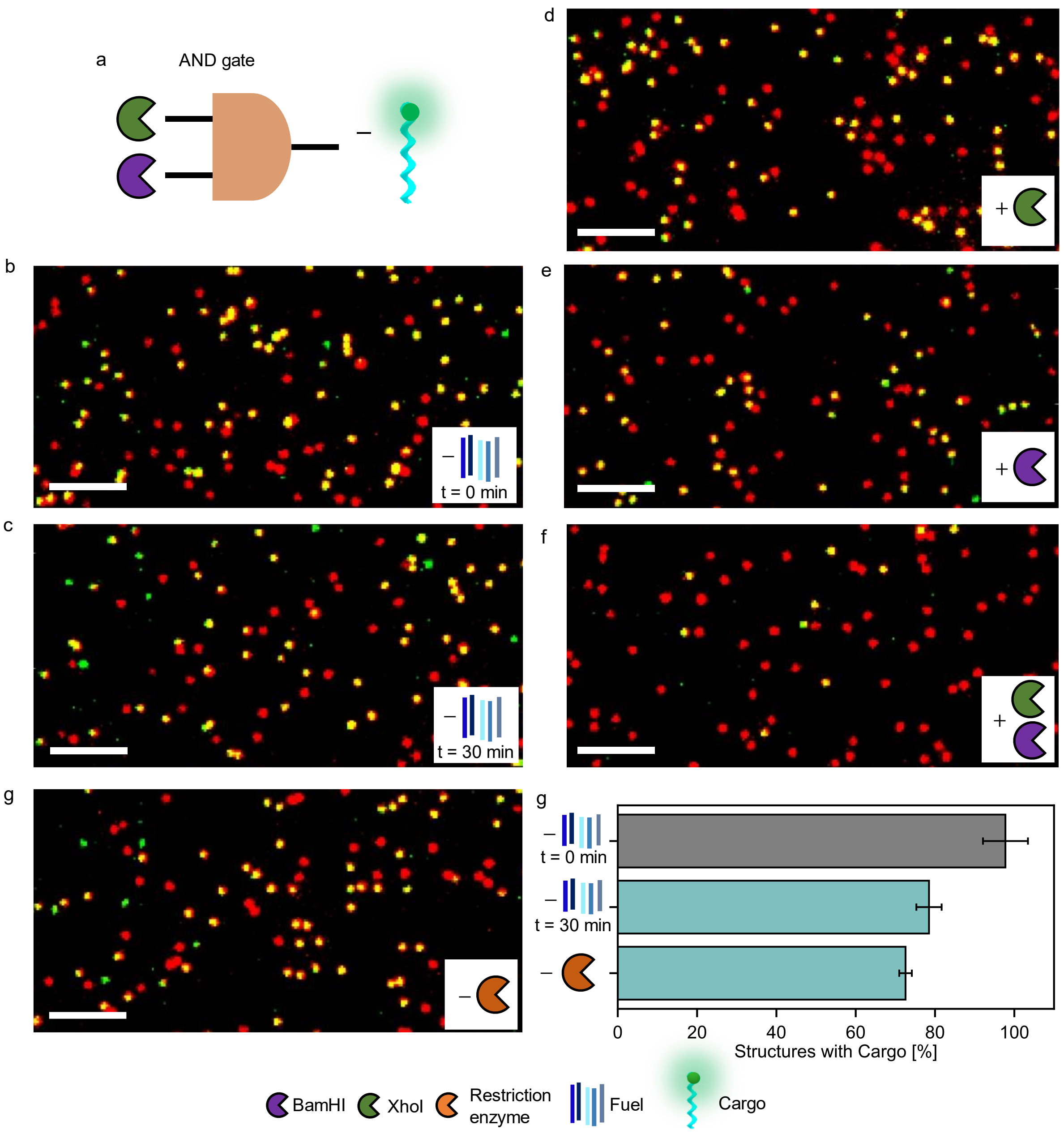


**Supplementary Figure 15. DNA origami arrays with AND logic gate releasing a cargo DNA strand in response to the combination of different restriction enzymes.** (a) Schematic representation of an AND logic gate which releases a cargo DNA strand in response to the activity of BamHI and XhoI. (b-f) Exemplary TIRF images of DNA origami array structures before and after incubation with different inputs. (g) Fraction of structures with cargo before and after incubation with and without fuel DNA strand but without restriction enzymes. In addition to possible inputs, all DNA origami arrays are incubated with fuel DNA strands 1-5. Error bars represent the standard deviation in the fractions calculated from three TIRF images. Scalebar: 4 µm.

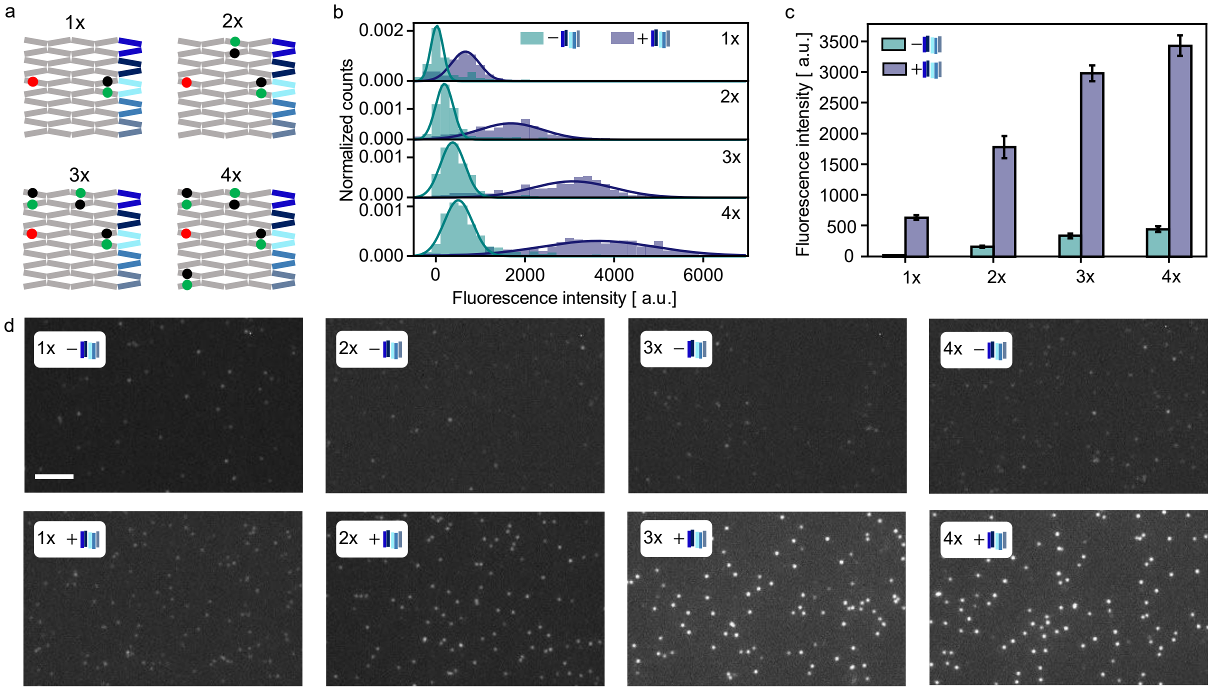


**Supplementary Figure 16. Signal amplification by incorporating multiple fluorescence onset output units into different antijunction nodes.** (a) DNA array designs for incorporating one, two, three and four green fluorescence onset units. Additionally, an ATTO647N dye is incorporated for locating the DNA array positions. ATTO542, BHQ2 (green onset unit) and ATTO647N are shown as green, black and red dots. (b) Fluorescence intensity distributions of ATTO542 for the different structures prior and after incubation with fuel DNA strands. When increasing the number of incorporated fluorescence onset units, the intensity contrast between the two states increases. Fluorescence intensities are calculated from dual-color TIRF images using ATTO647N fluorescence to locate the positions of the structures. The distributions are fitted with a Gaussian. (c) Mean values of the fluorescence intensity distributions for the different structures. Error bars represent the standard deviation in the mean values calculated from at least three TIRF images. (d) Exemplary TIRF images (ATTO542 excitation) for the DNA origami array structures shown in (a) before (upper row) and after (lower row) incubation with fuel DNA strands 1-5. Scalebar: 4 µm.

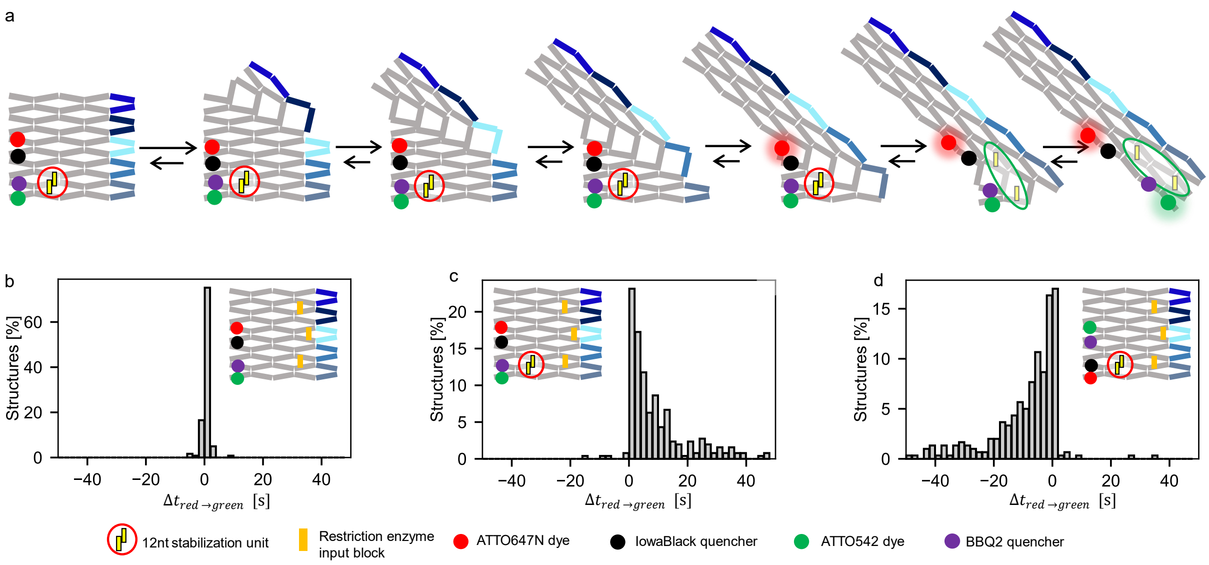


**Supplementary Figure 17. Controlling the timing and the order between output operations.** (a) A timing unit consisting of a 12 nt DNA lock is placed on antijunctions transforming before two red and green fluorescence onset units. (b-d) Distributions of the time difference between the fluorescence onset at the studied positions for a DNA origami array with BamHI-responsive input units (positions 1.1, 1.2, 1.3) obtained upon incubation with BamHI for DNA origamis (b) without and (c,d) with a timing unit incorporated. (b) Without the timing unit the red and green fluorescence onsets occur simultaneously. (c,d) with the timing unit, the red and green fluorescence onsets occur with a time delay between them. The order of fluorescence onsets hereby is controlled by the placement of the onset units with respect to the timing unit. In addition to BamHI, all DNA origami arrays are incubated with fuel DNA strands 1-5.

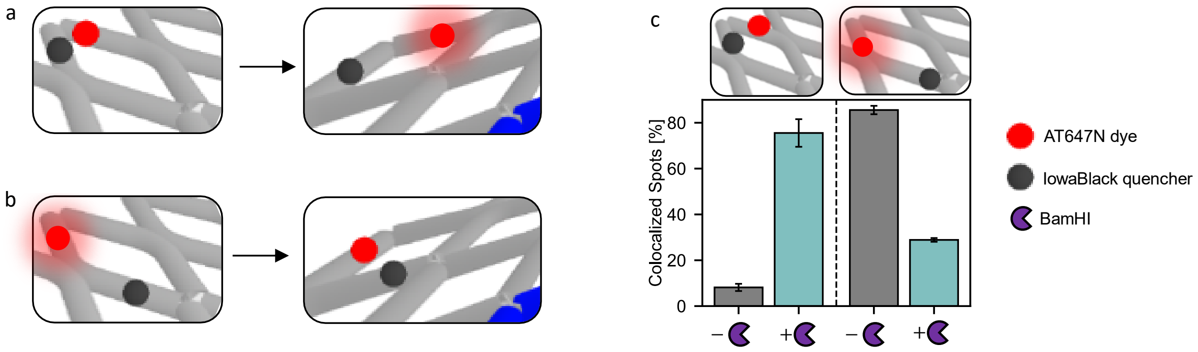


**Supplementary Figure 18. Concept of fluorescence onset and offset unit.** (a,b) Sketch demonstrating the principle of a fluorescence onset (a) and a fluorescent offset (b) unit. (c) Fraction of red-green colocalized spots on dual-color TIRF images of DNA origami arrays with a fluorescence on-/offset unit and three restriction enzyme locking units responsive to BamHI incorporated before and upon incubation with BamHI. In addition to BamHI, all DNA origami arrays are incubated with fuel DNA strands 1-5. Error bars represent the standard deviation in the fractions calculated from three TIRF images.

**Supplementary Note 3. Sequence of the p1800 scaffold used to fold the DNA origami array structure from 5' to 3' end:**

TACGAAGAGTTCCAGCAGGGATTCCAAGAAATGGCCAATGAAGATTGGATCACC
TTTCGCACTAAGACCTACTTGTTTGAGGAGTGCCTGATGAATTGGCACGACCGC
CTCAGGAAAGTGGAGGAGCATTCTGTGATGACTGTCAAGCTCCAATCTGAGGTG
GGCAAATATAAGATTGTTATCCCTATCTAGAAGTACGTCCGCGGAGAACACCTG
CCACCCGATCACTGGCTGGATCTGTTACGCTTGCTGGGTCTGCCTCGCGGCAC
ATCTCTGGAGAAACTGCTGTTCGGTGACCTGCTGAGAGTTGCCGATACCATCGT
GGCCAAGGCTGCTAACCTGAAAGATCTGAACTCACGCGGCCAGGGTGAAGTGA
CCATCCGCGAATAACTCAGGGAACTGGATTTGTGGGGCGTGGGTGCTGTGTTC
ACACTGATCGGCTATGAGGACTCCCAGAGCCGCACCTAGAAGCTGATCAAGGA
TTGGAAGGAGCTCGTCAACCAGGTGGGCGACAATATATGCCTCCTGCAGTCCTT
GAAGGACTCACCATACTATAAAGGCTTTGAAGACAAGGTCAGCATCTGGGCAAG
GAAACTCGCCGAACTGGACGATAATTTGCAGAACCTCAACCATATTCGCAGAAA
GTGGGTTTACCTCGAACCATACTTTGGTCGCGGAGCCCTGCCCAAAGAGCAGA
CCAGATTCAACAGGGTGGGCGAAGATTTCCGCAGCATCATGACATATATCAAGA
AGGACAATCGCGTCACGCCCTTGACTACCCACGCAGGCATTCTAAACTCACTGC
TGACCATCCTGGACCAATTGCAGAGATGCCAGCGCAGCCTCAACGAGTTCCTG
GAGGCGAAGCGCAGCGCCTTCCCTCGCTTTAACTTCATCGGAGACGATGACCT
GCGCGAGATCTTGGGCCAGTCAACCAATTAATCCGTGATTCAGTCTCACCTCAA
GAAGCTGTTTGCTGGTATCAACTCTGGCTGTTTCGATGAGAAGTCTAAGCACTAT
ACTGCAATGAAGTCCTTGGAGGGGCAAGTTGTGCCATTCAAGAATAACGTACCC
TTGTCCAATAACGTCGAAACCTGGCTGAACGATCTGGCCCTGGAGATGAAGAAG
ACCCTGGAGGCGCTGCTGAAGGAGTGCGTGACAACTAGACGCAGCTCTCAGGG
AGCTGTGGGCCCTTCTCTGTTCCCATCACAGATCTAGTGCTTGGCCGAACAGAT
CAAGTTTACCGAAGATGTGGAGAACGCAATTAAAGATCACTCCCTGCACCAGAT
TGAGTAACAGCTGGTGAACAAATTGGAGCAGTATACTAACATCGACACATCTTC
CGTAGACCCAGGTAACACAGAGTCCGGTATTCTGGAGCTGAAACTGAAAGCACT
GATTCTCGACGGATCCACGCGCCCTGTAGCGGCGCATTAAGCGCGGCGGGTGT
GGTGGTTACGCGCAGCGTGACCGCTACACTTGCCAGCGCCCTAGCGCCCGCTC
CTTTCGCTTTCTTCCCTTCCTTTCTCGCCACGTTCGCCGGCTTTCCCCGTCAAGC
TCTAAATCGGGGGCTCCCTTTAGGGTTCCGATTTAGTGCTTTACGGCACCTCGA
CCCCAAAAAACTTGATTTGGGTGATGGTTCACGTAGTGGGCCATCGCCCTGATA
GACGGTTTTTCGCCCTTTGACGTTGGAGTCCACGTTCTTTAATAGTGGACTCTTG
TTCCAAACTGGAACAACACTCAACCCTATCTCGGGCTATTCTTTTGATTTATAAG
GGATTTTGCCGATTTCGGGGTACC

**Table 1. Core staples from the 5‘ to the 3’ end for the DNA origami array structure.**

| ID | Sequence (5‘ to 3‘) | Function |
|---|---|---|
| C1 | GGAGTCCTCATAGATGGTTCGAGGTAAACCCACTTTCTGATGT CATGATGCT | core |
| C2 | GCGGAAATCTTCGAATTGGTTGACTGG CCCAAGATCTCGAGTGCTTAGACTT | core |
| C3 | GTTGAGGCTGCGCGGTCCAGGATGGTCAGCAGTGAGTTTCCC AGATGCTGAC | core |
| C4 | CTTGTCTTCAAAGAGTCCTTCAAGGACTGCAGGAGGCATCAAC TCTCAGCAG | core |
| C5 | AATCTTATATTTGATTCGCGGATGGTCACTTCACCCTGGAGGT GCGGCTCTG | core |
| C6 | GAGAGCTGCGTCTCCCCTCCAAG GACTTCATTGCAGTATCGCAGGT CATCGT | core |
| C7 | AGTGATCGGGTGGAGCAGCCTTGGCCACGATGGTATCGGATA TTGTCGCCCA | core |
| C8 | CCTGGTTGACGAGATCGTCCAGTTCGGCGAGTTTCCTTGAGA ATGCCTGCGT | core |
| C9 | CTCATCGAAACAGAGATCTGTGATGGGAACAGAGAAGGGACT CAATCTGGTGCAGGGAGTGATCTTTTTT | core |
| C10 | TTTTTGCGAAAGGTGATCCAATCTTCATTGGAACAGATCCAGC C | core |
| C11 | ATGGCACAACTTGAGTTGTCACGCACTCCTTCAGCAGCGACG GAAGATGTGTCGATGTTAGTATATTTTT | core |
| C12 | CTTCATCTCCAGGAGGTTTCGACGTTATTGGACAAGGGTCCTC CAGGAACTC | core |
| C13 | GTCACCGAACAGCTGCCGCGAGGCAGACCCAGCAAGCGTCC ATTTCTTGGAATCCCTGCTGGAAC | core |
| C14 | GGTTCTGCAAATTCTCCTTCCAATCCTTGATCAGCTTCTCCGC GTGAGTTCA | core |
| C15 | GATCTTTCAGGTTCAGGTGTTCTCCGCGGACGTACTTCTCACT CCTCAAACAAGTAGGTCTTAGTTTTTT | core |
| C16 | TTTTTTGAGGCGGTCGTGCCAATTCATCAGGAGATAGGGATAA C | core |
| C17 | TACCGGACTCTGTGTTACCTGGGTCTCCTCCAGGGTCTT | core |
| C18 | TTTTTCTGCTCCAATTTGTTCACCAGCTGTTCCCACAGCTCCCT | core |
| C19 | CTCCGATGAAGTTCGTGACGCGATTGTCCTTCTTGATATCGAA TATGGTTGA | core |
| C20 | GGGTAGTCAAGGGAAAGCGAGGGAAGGCGCTGCGCTTCGAC GTTATTCTTGA | core |
| X1 | CCAAATCAAGTTTTTTGGGGTCGAGGTGCCGTAAAGCACTAA | loop |
| X2 | AAACCGTCTATCAGGGCGATGGCCCACTACGTGAACCATCAC | loop |
| X3 | GTCCACTATTAAAGAACGTGGACTCCAACGTCAAAGGGCGAA | loop |
| X4 | GGTACCCCGAAATCGGCAAAATCCCTTATAAATCAAAAGAAT | loop |
| X5 | AAGGAGCGGGCGCTAGGGCGCTGGCAAGTGTAGCGGTCACG C | loop |
| X6 | AGCCCGAGATAGGGTTGAGTGTTGTTCCAGTTTGGAACAAGA | loop |

| | | |
|---|---|---|
| X7 | ATCGGAACCCTAAAGGGAGCCCCCGATTTAGAGCTTGACGGG | loop |
| X8 | TGCGCGTAACCACCACACCCGCCGCGCTTAATGCGCCGCTAC | loop |
| X9 | AGGGCGCGTGGATCCGTCGAGAATCAGTGCTTTCAGTTTCAG | loop |
| X10 | GAAAGCCGGCGAACGTGGCGAGAAAGGAAGGGAAGAAAGCG A | loop |

**Table 2. Fuel staples from the 5‘ to the 3’ end used to transform the DNA origami array structure.** Fuel staples are labelled according to the position they can bind to the DNA origami array structure, starting from the upper right corner to the lower right corner.

| **ID** | **Sequence (5‘ to 3‘)** | **Function** |
|---|---|---|
| T1 | AGTTCCCTGAGTTCCCACCTCAGATTGGAGCTTGACAGTCATC ACAGAAGCTCCTCCACTTTCCTTTTT | fuel 1 |
| T2 | CCGCGACCAAAGTCCGATCAGTGTGAACACAGCACCCACGCC CCACAAACC | fuel 2 |
| T3 | TGAATCACGGATTCCCACCCTGTTGAATCTGGTCTGCTCTTTG GGCAGGGT | fuel 3 |
| T4 | TCGGCCAAGCACTCCAGAGTTGATACCAGCAAACAGCTTCTTG AGGTGAGAC | fuel 4 |
| T5 | TTTTTTTAATTGCGTTCTCCACATCTTCGGTAAACTTGATCTGT | fuel 5 |
| T3* | TCTGGTCTGCTCTTTGGGCAGGGCT | fuel 3 - 25nt |

**Table 3. Staples from the 5‘ to the 3’ end for labelling the DNA origami array structure with biotin and fluorescent dye – quencher pairs.** B1 is included in all DNA array structures for surface immobilization. For the upper and lower fluorescence onset units (U and D staples) either the green or the red dye-quencher pair is used. The staples of the lower fluorescence onset unit (D1-D6) are used to incorporate AT542 and AT647N dyes for co-localization in the DNA arrays. Here, the corresponding quencher-labeled oligonucleotides (D4/ D6) are replaced by their unlabeled alternatives. For the red fluorescence offset unit O2 is used in combination with the staples of the left fluorescence onset unit (L1-L4). Here, the quencher labelled oligonucleotide L4 is replaced by the corresponding unlabeled strand.

| ID | Sequence (5‘ to 3‘) | Function | Replaces |
|---|---|---|---|
| B1 | **Biotin**-GAAAGCCGGCGAACGTGGCGAGAAAGGAAGGGAAGAAAGCGA | biotin | X10 |
| L1 | CTGCAGGAGGCATCAACTCTCAGCAG | onset/ offset Left replacement | C3, C4 |
| L2 | CAGTGAGTTTCCCAGATGCT | onset/ offset Left replacement | |
| L3 | GTTGAGGCTGCGCGGTCCAGGATGGTCAG-**ATTO647N** | onset/ offset Left red AT647N | |
| L4 | **IowaBlack**-GACCTTGTCTTCAAAGAGTCCTTCAAGGA | onset Left red Iowa Black | |
| U1 | CAGCAAGCGTCCATTTCTTG | onset Up replacement | C13 |
| U2 | **ATTO647N**-GAATCCCTGCTGGAAC | onset Up red AT647N | |
| U3 | GTCACCGAACAGCTGCCGCGAGGCAGACC-**IowaBlack** | onset Up red Iowa Black | |
| U4 | **ATTO542**-GAATCCCTGCTGGAAC | onset Up green AT542 | |
| U5 | GTCACCGAACAGCTGCCGCGAGGCAGACC-**BHQ2** | onset Up green BHQ2 | |
| D1 | ATTGGACAAGGGTCCTCCAGGAACTC | onset Low replacement | C12, C17 |
| D2 | ACCTGGGTCTCCTCCAGGGT | onset Low replacement | |
| D3 | **ATTO647N**-CTTCTTCATCTCCAGGAGGTTTCGACGTT | onset Low red AT647N | |
| D4 | TACCGGACTCTGTGTT-**IowaBlack** | onset Low red Iowa Black | |
| D5 | **ATTO542**-CTTCTTCATCTCCAGGAGGTTTCGACGTT | onset Low green AT542 | |
| D6 | TACCGGACTCTGTGTT-**BHQ2** | onset Low green BHQ2 | |
| O1 | CCTGGTTGACGAGATCGTCCAGTTCGGCGAGTTTCCTTGAGAATGCCTG-**IowaBlack** | offset Left red Iowa Black | C8 |
| Ri1 | GCCCAAGATCTCGAGTGCTTAGACTT | onset Right replacement | C1, C2 |
| Ri2 | CCACTTTCTGATGTCATGAT | onset Right replacement | |
| Ri3 | **ATTO542**-GCTGCGGAAATCTTCGAATTGGTTGACTG | onset Right green AT542 | |
| Ri4 | GGAGTCCTCATAGATGGTTCGAGGTAAAC-**BHQ2** | onset Right green BHQ2 | |
| URi 1 | CGTACTTCTCACTCCTCA | onset UpRight replacement | C15 |
| URi 2 | **ATTO542**-AACAAGTAGGTCTTAGT | onset UpRight green AT542 | |
| URi 3 | GATCTTTCAGGTTCAGGTGTTCTCCGCGGA-**BHQ2** | onset UpRight green BHQ2 | |

**Table 4. Staples from the 5' to the 3' end for incorporation of activation input units responsive to restriction enzyme activity and light.** Staples used to form locking units responsive to restriction enzyme activity (E-staples) and light of 300-350 nm (Li-staples). The part of the staples which is used to anchor them into the core structure is shown in black whereas linker sections added for flexibility and the stems which form the DNA locks are shown in blue and red, respectively. For E-staples, the part of the stem which is designed to be responsive to restriction enzyme activity is highlighted with six bold red Xs. This section is to be replaced by the cutting sequence of the restriction enzyme the corresponding input unit is designed to be responsive to (see Table 5). In Li-1, a spacer photocleavable by light of 300-350 nm is included after the linker sequence as denoted in the sequence by PC in purple.

| ID | Sequence (5' to 3') | Function | Replaces |
|---|---|---|---|
| E1 | TGAGGCGGTCGTGCCAATTCATCAGGAGATAGGGATAA TTTT GCCT **XXXXXX** GTGATGTAGGTGGTAGAGG | 1.1 Enzyme Unit 3' | C1, C2, C5, C9, C16 |
| E2 | CCTCTACCACCTACATCAC **XXXXXX** AGGC TTTT CACCCTGGAGGTGCGGCTCT | 1.1 Enzyme Unit 5' | |
| E3 | GGGAGTCCTCATAGATGGTTCGAG TTTT GCCT **XXXXXX** GTGATGTAGGTGGTAGAGG | 1.2 Enzyme Unit 3' | |
| E4 | CCTCTACCACCTACATCAC **XXXXXX** AGGC TTTT GCGGAAATCTTCGAATTGGTTGACTGGC | 1.2 Enzyme Unit 5' | |
| E5 | CCAAGATCTCGAGTGCTTAGACT TTTT GCCT **XXXXXX** GTGATGTAGGTGGTAGAGG | 1.3 Enzyme Unit 3' | |
| E6 | CCTCTACCACCTACATCAC **XXXXXX** AGGC TTTT GAGAAGGGACTCAATCTGGTGCAGGGAGTGATCT | 1.3 Enzyme Unit 5' | |
| E7 | TCTCATCGAAACAGAGATCTGTGATGGGAACA | 1.1-3 Enzyme Units Replacement | |
| E8 | GTAAACCCACTTTCTGATGTCATGATGCT | 1.1-3 Enzyme Units Replacement | |
| E9 | CAATCTTATATTTGATTCGCGGATGGTCACTT | 1.1-3 Enzyme Units Replacement | |
| E10 | TTGATCAGCTTCTCCGCGTGAGTTCAG TTTT GCCT **XXXXXX** GTGATGTAGGTGGTAGAGG | 2.1 Enzyme Unit 3' | C6, C14, C15, C18, C19 |
| E11 | CCTCTACCACCTACATCAC **XXXXXX** AGGC TTTT GACGTACTTCTCACTCCTCAAACAAGTAGGTCTTAGT | 2.1 Enzyme Unit 5' | |
| E12 | CTCCGATGAAGTTCGTGACGCGATTGTCCTTT TTTT GCCT **XXXXXX** GTGATGTAGGTGGTAGAGG | 2.2 Enzyme Unit 3' | |

| | | | |
|---|---|---|---|
| E13 | CCTCTACCACCTACATCAC **XXXXXX** AGGC TTTT TTTTCTGCAAATTCTCCTTCCAATCC | 2.2 Enzyme Unit 5' | |
| E14 | CTGCTCCAATTTGTTCACCAGCTGTTCCCACAGCTC CCTG TTTT GCCT **XXXXXX** GTGATGTAGGTGGTAGAGG | 2.3 Enzyme Unit 3' | |
| E15 | CCTCTACCACCTACATCAC **XXXXXX** AGGC TTTT CATTGCAGTATCGCAGGTCATCGT | 2.3 Enzyme Unit 5' | |
| E16 | AGAGCTGCGTCTCCCCTCCAAGGACTT | 2.1-3 Enzyme Units Replacement | |
| E17 | TCTTGATATCGAATATGGTTGAGG | 2.1-3 Enzyme Units Replacement | |
| E18 | ATCTTTCAGGTTCAGGTGTTCTCCGCG | 2.1-3 Enzyme Units Replacement | |
| E19 | GCGAAAGGTGATCCAATCTTCATTGGAACAGATCCA G TTTTTT GCCT **XXXXXX** GTGATGTAGGTGGTAGAGG | 3.1 Enzyme Unit 3' | C7, C8, C10, C11, C20 |
| E20 | CCTCTACCACCTACATCAC **XXXXXX** AGGC TTTTTT GCCACGATGGTATCGGATATTGTCGCC | 3.1 Enzyme Unit 5' | |
| E21 | CACCTGGTTGACGAGATCGTCCAGTT TTTTTT GCCT **XXXXXX** GTGATGTAGGTGGTAGAGG | 3.2 Enzyme Unit 3' | |
| E22 | CCTCTACCACCTACATCAC **XXXXXX** AGGC TTTTTT CGTGGGTAGTCAAGGGAAAGCGAGGGAAG | 3.2 Enzyme Unit 5' | |
| E23 | GCGCTGCGCTTCGACGTTATTCTT TTTTTT GCCT **XXXXXX** GTGATGTAGGTGGTAGAGG | 3.3 Enzyme Unit 3' | |
| E24 | CCTCTACCACCTACATCAC **XXXXXX** AGGC TTTTTT CACTCCTTCAGCAGCGACGGAAGATGTGTCGATGTT AGTATA | 3.3 Enzyme Unit 5' | |
| E25 | CCAGTGATCGGGTGGAGCAGCCTTG | 3.1-3 Enzyme Units Replacement | |
| E26 | CGGCGAGTTTCCTTGAGAATGCCTG | 3.1-3 Enzyme Units Replacement | |
| E27 | GAATGGCACAACTTGAGTTGTCACG | 3.1-3 Enzyme | |

| | | | |
|---|---|---|---|
| | | Units Replacement | |
| Li1 | TGAGGCGGTCGTGCCAATTCATCAGGAGATAGGGATAA TTTT **PC** GCCTAAGCTTGTGATGTAGGTGGTAGAGG | 1.1 Light Unit 3' | E1, E2 |
| Li2 | CCTCTACCACCTACATCACAAGCTTAGGC TTTT CACCCTGGAGGTGCGGCTCT | 1.1 Light Unit 5' | |

**Table 5. Cutting sites of the used restriction enzymes from the 5‘ to the 3’ end.** Sequences for the cut sites of the restriction enzymes BamHI, XhoI and StuI. Additionally, a fourth sequence is shown which is not cleavable by any of the aforementioned enzymes. Depending on which enzyme an input unit is designed to be responsive to, the sequences replace the red Xs for the corresponding E-staples in Table 4.

| Restriction enzyme | Cut site (5‘ to 3‘) |
|---|---|
| BamHI | GGATCC |
| XhoI | CTCGAG |
| StuI | AGGCCT |
| not activatable | AAGCTT |

**Table 6. Staples from the 5‘ to the 3’ end for incorporation of an inhibition input units responsive to the anti-Dig antibody.**

| ID | Sequence (5‘ to 3‘) | Function | Replaces |
|---|---|---|---|
| A1 | TGAGGCGGTCGTGCCAATTCATCAGGAGATAGGGATAAC-**Dig** | Antibody Unit - Antigen 1 | C16 |
| A2 | TAACATTCCTAACTTCTCATACTCATCGAAACAGAGATCTGTG | Antibody Unit - Antigen 2 site | C9 |
| A3 | ATGGGAACAGAGAAGGACTCAATCTGGTG | Antibody Unit - Antigen 2 Replacement | |
| A4 | **Dig**-TTATGAGAAGTTAGGAATGTTA | Antibody Unit - Antigen 2 | |

**Table 7. Staples from the 5‘ to the 3’ end for incorporation of activation and inhibition input units responsive to a 20 nt ssDNA input.** The part of the staples which is used to anchor them into the core structure is shown in black. Linker sections added for flexibility and the ssDNA binding sites with toehold at the 5’ are shown in blue and red, respectively.

| ID | Sequence (5‘ to 3‘) | Function | Replaces |
|---|---|---|---|
| DNA1 | TGAGGCGGTCGTGCCAATTCATCAGGAGA TAGGGATAA TTT GCTCGACTGATG | 1.1 DNA Activation/Inhibition Unit 3' | C1, C2, C5, C9, C16 |
| DNA2 | TGCAGTCG CATCAGTCGAGC TTT CACCCTGGAGGTGCGGCTCT | 1.1 DNA Activation Unit 5' | |
| DNA3 | GGGAGTCCTCATAGATGGTTCGAG TTT GCTCGACTGATG | 1.2 DNA Activation/ Inhibition Unit 3' | |
| DNA4 | TGCAGTCG CATCAGTCGAGC TTT GCGGAAATCTTCGAATTGGTTGACTGGC | 1.2 DNA Activation Unit 5' | |
| DNA5 | CCAAGATCTCGAGTGCTTAGACT TTT GCTCGACTGATG | 1.3 DNA Activation/ Inhibition Unit 3' | |
| DNA6 | TGCAGTCG CATCAGTCGAGC TTT GAGAAGGGACTCAATCTGGTGCAGGGAGT GATCT | 1.3 DNA Activation Unit 5' | |
| DNA7 | AACCCACTTTCTGATGTCATGAT | 1.1-3 DNA Units Replacement | |
| DNA8 | TCTTATATTTGATTCGCGGATGGTCA | 1.1-3 DNA Units Replacement | |
| DNA9 | CATCGAAACAGAGATCTGTGATGGGA | 1.1-3 DNA Units Replacement | |
| DNA10 | GCGAAAGGTGATCCAATCTTCATTGGAACA GATCCAG TTT GCTCGACTGATG | 3.1 DNA Activation Unit 3' | C7, C8, C10, C11, C20 |
| DNA11 | TGCAGTCG CATCAGTCGAGC TTT GTATCGGATATTGTCGCC | 3.1 DNA Activation Unit 5' | |
| DNA12 | CACCTGGTTGACGAGATCGTCCAGTT TTT GCTCGACTGATG | 3.2 DNA Activation Unit 3' | |
| DNA13 | TGCAGTCG CATCAGTCGAGC TTT TGGGTAGTCAAGGGAAAGCGAGGGAAG | 3.2 DNA Activation Unit 5' | |
| DNA14 | GCGCTGCGCTTCGACGTTATTCTT TTT GCTCGACTGATG | 3.3 DNA Activation Unit 3' | |
| DNA15 | TGCAGTCG CATCAGTCGAGC TTT AGCAGCGACGGAAGATGTGTCGATGTTAG TATA | 3.3 DNA Activation Unit 5' | |
| DNA16 | GTGATCGGGTGGAGCAGCCTTGGCCACG | 3.1-3 DNA Units Replacement | |
| DNA17 | TGGCACAACTTGAGTTGTCACGCACTCC | 3.1-3 DNA Units Replacement | |
| DNA18 | CGAGTTTCCTTGAGAATGCCT | 3.1-3 DNA Units Replacement | |
| DNA19 | TGCAGTCG CATCAGTCGAGC TTT GACGTACTTCTCACTCCTCAAACAAGTAGG TCTTAGT | 1.1 DNA Inhibition Unit 5' | E11, E13, E15 |
| DNA20 | TGCAGTCG CATCAGTCGAGC TTT TTTTCTGCAAATTCTCCTTCCAATCC | 1.2 DNA Inhibition Unit 5' | |

| | | | |
|---|---|---|---|
| DNA21 | TGCAGTCG CATCAGTCGAGC TTT CATTGCAGTATCGCAGGTCATCGT | 1.3 DNA Inhibition Unit 5' | |
| DNA22 | TGAGGCGGTCGTGCCAATTCATCAGGAGA TAGGGATAA TTT GTGATGTAGGTG | 1.1 DNA Destabilization Unit 3' | DNA1 |
| DNA23 | CACCTACATCAC TTT GACGTACTTCTCACTCCTCAAACAAGTAGG TCTTAGT | 1.2 DNA Destabilization Unit 5' | DNA19 |
| DNA24 | GACTGATG CGACTG | 14nt ssDNA input | - |

**Table 8. Staples from the 5‘ to the 3’ end for incorporation of a cargo release output operation unit.** The part of the staples which is used to anchor them into the core structure is shown in black whereas linker sections added for flexibility and the stems which form the DNA locks are shown in blue and red, respectively. The part of the sequence to which the ATTO542-labelled cargo ssDNA strand which is to be released is hybridized is shown in purple.

| ID | Sequence (5‘ to 3‘) | Function | Replaces |
|---|---|---|---|
| R1 | TCCTCTACCA GTATCGTAG TTTTTTTTTT AGCAGTGAGTTTCCCAGATGCTGACCT | Cargo release - Catching unit 5' | C3, C4 |
| R2 | TGTCTTCAAAGAGTCCTTCAAGGACTGC TTTTTTTTTT CTACGATAC CCTACATCAC | Cargo release - Catching unit 3' | |
| R3 | AGGAGGCATCAACTCTCAGCAG | Cargo release - Replacement | |
| R4 | GTTGAGGCTGCGCGGTCCAGGATGGTC | Cargo release - Replacement | |
| R5 | GTGATGTAGGTGGTAGAGGAT-**ATTO542** | Cargo strand | - |

**Table 9. Staples from the 5‘ to the 3’ end for incorporation of the timing unit used to retard the transformation at a specific position.** The part of the staples which is used to anker them into the core structure is shown in black. Linker sections added for flexibility and the 12 nt long locking unit which forms a stem are shown in blue and red, respectively.

| ID | Sequence (5‘ to 3‘) | Function | Replaces |
|---|---|---|---|
| T1 | GGGTAGTCAAGGGAAAGCGAGGGAAGGCGCTG CGCTTCGACGTTATTCTTGAATG TTT CGACTACGATAC | Timing Unit - Locking unit 12bp | C11, C20 |
| T2 | GCACAACTTGAGTTGTCACGCACTCCTTCAGCA GCGACGGAAGATGTGTCGA T GTATCGTAGTCG | Timing Unit - Locking unit 12bp | |

**Table 10. DNA origami array designs used in this work.** For each DNA origami design, the modified staples are listed. Staples which are used as given in Tables 3-9 are listed without additional notation. Staples which are used without their corresponding modification (either highlighted in color or in bold in Tables 3-9) are listed with the addition _um. For E-staples, the included cutting site of the corresponding enzyme (Table 5) is highlighted by the addition _Enzyme or _nonAct for the corresponding enzyme cutting site and the non-cleavable sequence, respectively. The modified staples replace staples of the core mix (C- and L-staples as highlighted in Tables 3-9). For DNA origami folding, in addition to the modified staples, all unreplaced core staples are added.

| No | Modified Staples | Description |
|---|---|---|
| 1 | B1, L1-4, D1-2, D5, D6_um, DNA1-9 | DNA Activation Block Introduction (3x Block in Col1, Fig. 2) |
| 2 | B1, L1-4, D1-2, D5, D6_um, DNA1, DNA3, DNA5, DNA 7-9, DNA19-21, DNA2_um, DNA4_um, DNA6_um, E10_um, E12_um, E14_um, E16-18 | DNA Inhibition Block Introduction (3x Block in Col1,2, Fig. 2) |
| 3 | B1, L1-4, D1-2, D5, D6_um, DNA1-18 | 6x DNA Activation Block (Fig. 2) |
| 4 | B1, L1-4, D1-2, D5, D6_um, E1-4_XhoI, E5-6_um, E7-9 | XhoI Block Col1 Up, Mid (design 1, Fig. 3) |
| 5 | B1, L1-4, D1-2, D5, D6_um, E1-2_um, E3-6_XhoI, E7-9 | XhoI Block Col1 Mid, Low (design 2, Fig. 3) |
| 6 | B1, L1-4, D1-2, D5, D6_um, E1-6_StuI, E7-9, E10-15_XhoI, E16-18 | AND gate (XhoI, StuI, Fig. 3) |
| 7 | B1, L1-4, D1-2, D5, D6_um, E1-2_StuI, E3-4_um, E5-6_XhoI, E7-9 | OR gate (XhoI, StuI, Fig. 3) |
| 8 | B1, L1-4, D1-2, D5, D6_um, Li1-2, E3-4_um, E5-6_XhoI, E7-9 | OR gate (XhoI, Light, Fig. 3) |
| 9 | B1, L1-4, D1-2, D5, D6_um, E1-2_nonAct, E3-4_StuI, E5-6_XhoI, E7-9, E19-24_BamHI, E25-27 | 3xAND gate (XhoI, StuI, BamHI, Fig. 3) |
| 10 | B1, D1-3, D4_um, E1-6_BamHI, E7-9, R1-5 | Cargo release by BamHI activity (Fig. 4) |
| 11 | B1, D1-3, D4_um, E1-2_nonAct, E3-4_BamHI, E5-6_XhoI, E7-9, R1-5 | Cargo release by AND gate (BamHI, XhoI, Fig. 4) |
| 12 | B1, D1-3, D4_um, E1-2_BamHI, E3-4_um, E5-6_XhoI, E7-9, R1-5 | Cargo release by OR gate (BamHI, XhoI, Fig. 4) |
| 13 | B1, L1-3, L4_um, O1, D1-2, D5-6, DNA1-6, E7-9, E10-15_XhoI, E16-18 | full nanorobot with $\Delta t = 0$ (Fig. 5) |
| 14 | B1, L1-3, L4_um, O1, D1-2, D5-6, DNA1-6, E7-9, E10-15_XhoI, E16-18, Ti1-2 | full nanorobot with $\Delta t > 0$ (Fig. 5) |
| 15 | B1, L1-4, D1-2, D5, D6_um, E1-6_BamHI, E7-9 | BamHI Block Introduction (Fig. S2) |
| 16 | B1, L1-4, D1-2, D5, D6_um, E1-6_XhoI, E7-9 | XhoI Block Introduction (Fig. S2) |
| 17 | B1, L1-4, D1-2, D5, D6_um, E1-6_StuI, E7-9 | StuI Block Introduction (Fig. S2) |
| 18 | B1, L1-4, D1-2, D5, D6_um, Li1-2, E3-4_um, E5-6_nonAct, E7-9 | Light Block Introduction (Fig. S3) |
| 19 | B1, L1-4, D1-2, D5, D6_um, A1-4 | Dig Block Introduction (Fig. S4) |
| 20 | B1, L1-4, D1-2, D5, D6_um, DNA3-18, DNA1-2_um | 5x DNA Activation Block (Fig. S6) |

| | | |
|---|---|---|
| 21 | B1, L1-4, D1-2, D5, D6_um, DNA5-18, DNA1-4_um | 4x DNA Activation Block (Fig. S6) |
| 22 | B1, L1-4, D1-2, D5, D6_um, DNA1, DNA3-19, DNA2_um, E11-15_um, E16-18 | 5x DNA Activation Block + 1x Inhibition Block (Fig. S6) |
| 23 | B1, L1-4, D1-2, D5, D6_um, DNA3-18, DNA22-23, DNA2_um, E11-15_um, E16-18 | 5x DNA Activation Block + 1x Destabilization Block (Fig. S6) |
| 24 | B1, L1-4, D1-2, D5, D6_um, E1-2_XhoI, E3-6_um, E7-9 | XhoI Block Col1 Up (Fig. S8) |
| 25 | B1, L1-4, D1-2, D5, D6_um, E1-2_um, E3-4_XhoI, E5-6_um, E7-9 | XhoI Block Col1 Mid (Fig. S8) |
| 26 | B1, L1-4, D1-2, D5, D6_um, E1-4_um, E5-6_XhoI, E7-9 | XhoI Block Col1 Low (Fig. S8) |
| 27 | B1, L1-4, D1-2, D5, D6_um, E1-2_XhoI, E4-5_um, E5-6_XhoI, E7-9 | XhoI Block Col1 Up, Low (Fig. S8) |
| 28 | B1, L1-4, D1-2, D5, D6_um, E1-6_XhoI, E7-9 | XhoI Block Col1 Up, Mid, Low (Fig. S8) |
| 29 | B1, L1-4, D1-2, D5, D6_um, E10-11_XhoI, E12-15_um, E16-18 | XhoI Block Col2 Up (Fig. S8) |
| 30 | B1, L1-4, D1-2, D5, D6_um, E10-11_um, E12-13_XhoI, E14-15_um, E16-18 | XhoI Block Col2 Mid (Fig. S8) |
| 31 | B1, L1-4, D1-2, D5, D6_um, E10-13_um, E14-15_XhoI, E16-18 | XhoI Block Col2 Low (Fig. S8) |
| 32 | B1, L1-4, D1-2, D5, D6_um, E10-13_XhoI, E14-15_um, E16-18 | XhoI Block Col2 Up, Mid (Fig. S8) |
| 33 | B1, L1-4, D1-2, D5, D6_um, E10-11_XhoI, E12-13_um, E14-15_XhoI, E16-18 | XhoI Block Col2 Up, Low (Fig. S8) |
| 34 | B1, L1-4, D1-2, D5, D6_um, E10-11_um, E12-15_XhoI, E16-18 | XhoI Block Col2 Mid, Low (Fig. S8) |
| 35 | B1, L1-4, D1-2, D5, D6_um, E10-15_XhoI, E16-18 | XhoI Block Col2 Up, Mid, Low (Fig. S8) |
| 36 | B1, L1-4, D1-2, D5, D6_um, E19-24_XhoI, E25-27 | XhoI Block Col3 Up, Mid, Low (Fig. S8) |
| 37 | B1, L1-4, D1-2, D5, D6_um, E1-2_XhoI, E3-6_um, E7-9, E19-24_XhoI, E25-27 | XhoI Block Col1 Up; Col3 Up, Mid, Low (Fig. S8) |
| 38 | B1, L1-4, D1-2, D5, D6_um, E1-2_um, E3-4_XhoI, E5-6_um, E7-9, E19-24_XhoI, E25-27 | XhoI Block Col1 Mid; Col3 Up, Mid, Low (Fig. S8) |
| 39 | B1, L1-4, D1-2, D5, D6_um, E1-4_um, E5-6_XhoI, E7-9, E19-24_XhoI, E25-27 | XhoI Block Col1 Low; Col3 Up, Mid, Low (Fig. S8) |
| 40 | B1, L1-4, D1-2, D5, D6_um, E1-4_XhoI, E5-6_um, E7-9, E19-24_XhoI, E25-27 | XhoI Block Col1 Up, Mid; Col3 Up, Mid, Low (Fig. S8) |
| 41 | B1, L1-4, D1-2, D5, D6_um, E1-2_um, E3-6_XhoI, E7-9, E19-24_XhoI, E25-27 | XhoI Block Col1 Mid, Low; Col3 Up, Mid, Low (Fig. S8) |
| 42 | B1, L1-4, D1-2, D5, D6_um, E1-2_XhoI, E4-5_um, E5-6_XhoI, E7-9, E19-24_XhoI, E25-27 | XhoI Block Col1 Up, Low; Col3 Up, Mid, Low (Fig. S8) |
| 43 | B1, L1-4, D1-2, D5, D6_um, E1-2_StuI, E3-4_um, E5-6_XhoI, E7-9, E19-24_BamHI, E25-27 | 2outOf3 gate (XhoI, StuI, BamHI, Fig. S11) |

| | | |
|---|---|---|
| 44 | B1, D1-2, D5-6, E1-6_StuI, E7-9 | StuI Block + green Onset Block (Multiplexing, Fig. S12) |
| 45 | B1, D1-4, E1-6_XhoI, E7-9 | XhoI Block + red Onset Block (Multiplexing, Fig. S12) |
| 46 | B1, L1-3, L4_um, Ri1-4 | 1x green Onset Block (Fig. S16) |
| 47 | B1, L1-3, L4_um, Ri1-4, URi1-3 | 2x green Onset Block (Fig. S16) |
| 48 | B1, L1-3, L4_um, Ri1-4, URi1-3, U1, U4-5 | 3x green Onset Block (Fig. S16) |
| 49 | B1, L1-3, L4_um, Ri1-4, URi1-3, U1, U4-5, D1-2, D5-6 | 4x green Onset Block (Fig. S16) |
| 50 | B1, L1-4, D1-2, D5-6, E1-6_BamHI, E7-9 | red Onset, green Onset simultaneous (Fig. S17) |
| 51 | B1, L1-4, D1-2, D5-6, E1-6_BamHI, E7-9, Ti1-2 | red Onset, green Onset delayed (Fig. S17) |
| 52 | B1, L1-2, L5-6, D1-4, E1-6_BamHI, E7-9, Ti1-2 | green Onset, red Onset delayed (Fig. S17) |
| 53 | B1, L1-3, L4_um, O1 D1-2, D5, D6_um, E1-6_BamHI, E7-9 | red Offset Block Introduction (+BamHI, Fig. S18) |

### Publication bibliography

Shaw, Alan; Hoffecker, Ian T.; Smyrlaki, Ioanna; Rosa, Joao; Grevys, Algirdas; Bratlie, Diane et al. (2019): Binding to nanopatterned antigens is dominated by the spatial tolerance of antibodies. In *Nature nanotechnology* 14 (2), pp. 184–190. DOI: 10.1038/s41565-018-0336-3.
